# The Metabesity Factor HMG20A Potentiates Astrocyte Survival and Reactivity Preserving Neuronal Integrity

**DOI:** 10.1101/2021.02.15.431213

**Authors:** Petra I. Lorenzo, Esther Fuente-Martín, José M. Mellado-Gil, José A. Guerrero Martínez, Nadia Cobo-Vuilleumier, Valentine Comaills, Eugenia Martin Vazquez, Silvana Y. Romero-Zerbo, Jaime Muñoz Franco, Jesús A. Perez-Cabello, Sabrina Rivero Canalejo, Antonio Campos-Caro, Christian Claude Lachaud, Manuel Aguilar-Diosdado, Eduardo García Fuentes, Alejandro Martin-Montalvo, Manuel Álvarez Dolado, Franz Martin, Gemma Rojo-Martinez, David Pozo, Francisco J. Bérmudez-Silva, José C. Reyes, Benoit R. Gauthier

**Affiliations:** Andalusian Center of Molecular Biology and Regenerative Medicine-CABIMER, Junta de Andalucía-University of Pablo de Olavide-University of Seville-CSIC, Seville, Spain; Unidad de Gestión Clínica Intercentros de Endocrinología y Nutrición, Instituto de Investigación Biomédica de Málaga (IBIMA), Hospital Regional Universitario de Málaga, Universidad de Málaga, Spain; Department of Normal and Pathological Histology and Cytology, University of Seville School of Medicine, Seville, Spain; University Hospital “Puerta del Mar”, Instituto de Investigación e Innovación en Ciencias Biomédicas de la Provincia de Cádiz (INiBICA), Cádiz, Spain; Endocrinology and Metabolism Department, University Hospital “Puerta del Mar”, Instituto de Investigación e Innovación en Ciencias Biomédicas de la Provincia de Cádiz (INiBICA), Cádiz, Spain; Unidad de Gestión Clínica de Aparato Digestivo, Hospital Universitario Virgen de la Victoria, Instituto de Investigación Biomédica de Málaga (IBIMA), Spain; Centro de Investigación Biomédica en Red de Diabetes y Enfermedades Metabólicas Asociadas (CIBERDEM), Madrid, Spain

**Keywords:** HMG20A, metabolism, metabesity, astrocytes, inflammation

## Abstract

**Rationale:** We recently demonstrated that the ‘Metabesity’ factor HMG20A regulates islet beta-cell functional maturity and adaptation to physiological stress such as pregnancy and pre-diabetes. HMG20A also dictates central nervous system (CNS) development via inhibition of the LSD1/CoREST complex but its expression pattern and function in adult brain remains unknown. Herein we sought to determine whether HMG20A is expressed in the adult CNS, specifically in hypothalamic astrocytes that are key in glucose homeostasis and whether similar to islets, HMG20A potentiates astrocyte function in response to environmental cues.

**Methods:** HMG20A expression profile was assessed by quantitative PCR (RT-PCR) and/or immunofluorescence in: 1) the hypothalamus of mice exposed or not to a high-fat diet, 2) human blood leukocytes and adipose tissue obtained from healthy or diabetic individuals 3) primary mouse hypothalamic astrocytes exposed to either high glucose or palmitate. To investigate the function and regulatory mechanism of HMG20A, RNA-seq and cell metabolic parameters were performed on astrocytes treated or not with a siHMG20A. The regulatory function of HMG20A on astrogliosis was also assessed pharmacologically using ORY1001. Astrocyte-mediated neuronal survival was evaluated using conditioned media from siHMG20A-treated astrocytes.

**Results:** We show that *Hmg20a* is predominantly expressed in hypothalamic astrocytes, the main nutrient-sensing cell type of the brain. *Hmg20A* expression was upregulated in diet-induced obesity and glucose intolerant mice, correlating with increased transcript levels of *Gfap* and *Il1b* indicative of inflammation and astrogliosis. Expression levels were also increased in adipose tissue of obese non-diabetic individuals as compared to obese diabetic patients. HMG20A silencing in astrocytes resulted in repression of inflammatory, cholesterol biogenesis and epithelial-to-mesenchymal transition pathways with a concomitant increase in apoptosis and reduced mitochondrial bioenergetics. Motoneuron viability was also hindered in HMG20A-depleted astrocyte-derived conditioned media. Astrogliosis was induced using ORY1001, a pharmacological inhibitor of the LSD1/CoREST complex, mimicking the effect of HMG20A.

**Conclusion:** HMG20A coordinates the astrocyte polarization state. Under physiological pressure such as obesity and insulin resistance that induces low grade inflammation, HMG20A expression is increased to induce astrogliosis in an attempt to preserve the neuronal network and glucose homeostasis. Nonetheless, a chronic metabesity state or functional mutations will result in lower levels of HMG20A, failure to promote astrogliosis and increase susceptibility of neurons to stress-mediated apoptosis. Such effects could be therapeutically reversed by ORY1001-induced astrogliosis.

## Introduction

‘Metabesity’ is an emerging global term describing a wide range of diseases with underlying metabolic disorders and for which the etiology lies in complex interactions among genes and the obesogenic environment [1, 2]. Metabesity contributes to the generation of a chronic low-grade inflammation [3], which is recognized as the pathogenic link with Type 2 Diabetes Mellitus (T2DM). This inflammation targets peripheral tissues specialized in metabolism such as liver, adipose tissue and pancreatic islets and plays a key role in insulin resistance and beta-cell dysfunction [4-6]. Inflammation in the central nervous system (CNS) is an early event detected even prior to weight gain, and is likely implicated in the increased incidence of neurodegenerative diseases among T2DM patients [7]. In fact, DM and neurodegenerative disease such as Alzheimer’s disease (AD), share common elements such as insulin resistance, altered glucose metabolism, amyloid aggregation and inflammatory response [8]. Accordingly, alterations in the insulin-central nervous system axis and brain glucose metabolism in DM and/or obese individual have been associated with the progression of AD [9, 10]. Of particular interest, hypothalamic astrocytes, the major glucose metabolizers in the brain, were shown to couple CNS glucose uptake and nutrient availability via insulin signalling [10]. Consequently, astrocytes within the hypothalamus, the hub of metabolic homeostasis, have come into the limelight as key regulator of appetite and energy expenditure.

Astrocytes are dispersed throughout the brain and regulate tissue homeostasis through structural and nutritive support of neurons, energy storage, and defense against oxidative stress. In view of their atypical morphology characterized by elongated processes of the cellular membrane (hence their name derived from star-like cells) that are in physical contact with other cell types and blood vessels, astrocytes facilitate synaptic transmission, neuronal plasticity and survival as well as to regulate blood flow relaying incoming peripheral signals to the CNS [3]. As such astrocytes express a multitude of receptors for hormones and neuropeptides involved in metabolic control and accordingly to the registered signal will react by modulating the neuronal circuitry through increased transport of metabolic factors and release of growth factors, gliotransmitters, metabolites and cytokines [11]. Notwithstanding to these physiological functions, astrocytes also react/adapt to CNS insults such as metabesity-induced neuroinflammation, through a complex and multifaceted process called astrogliosis, characterized by cellular hypertrophy, increased proliferation and altered astrocyte functions [12]. Although astrogliosis has an initial protective effect to preserve neuronal function and viability as well as repairing/healing of the initial damage, failure to resolve this reactive phenotype contributes to injury and to the development and progression of AD and DM [13, 14]. Indeed, astrogliosis impairs insulin signalling and glycogen storage, increases glucose uptake and glycolysis, and alters glucose transport in the brain [3]. Although well characterized at the cellular level, the molecular mechanism controlling astrogliosis remains ill-defined with only few factors such as ZEB2, an epithelial-to-mesenchymal transition (EMT)-related transcription factor or NF-kB, a pro-inflammatory transcriptional regulator which were found to promote astrogliosis [15, 16]. As such, identifying key regulators of astrocyte phenotypic state may prove valuable in the development of novel DM and AD theranostics.

Of particular interest is the chromatin remodeling factor HMG20A that we recently reported to be essential for pancreatic islet beta-cell functional maturation and adaptation to stress conditions such as hyperglycemia and pregnancy [17, 18]. Consistent with these functional data small nucleotide polymorphisms (SNPs) in the *HMG20A* gene have been associated to T2DM as well as gestational diabetes mellitus (GDM) in Asian/Indian and European populations [19-25]. HMG20A is also a key regulator of neuronal differentiation [26] by exerting global genomic changes through establishing active or silent chromatin [27]. Mechanistically, HMG20A competes with HMG20B to relieve the transcriptional repression imposed by the complex LSD1-CoREST histone demethylase, which function is to silence neuronal and islet beta-cell genes through its interaction with the transcription factor REST [28-31]. As HMG20A regulates the expression of key genes such as NEUROD, GK, and GLUTs common to both astrocytes and beta-cells we reason that this chromatin factor is a common master regulator of beta-cells and astrocytes, integrating inputs to altered glucose levels and stressful physiological conditions that dictate the phenotypic status of these cells [3]. As such, herein, we explored whether HMG20A is expressed in astrocytes and whether it potentiates astrocytes function and phenotypic state in response to stress conditions. We also assessed whether HMG20A is expressed in human white adipose tissue (WAT) and serum and whether it’s levels correlates with obesity and/or T2DM. Our studies identify HMG20A as a core regulator of molecular pathways implicated in astrogliosis a critical process for neuron survival in response to injury and/or stress. Accordingly, higher levels of HMG20A in WAT also correlated with a healthy status in obese individuals.

## Methods

### Ethical Statement

The collection of human samples was carried out in accordance with the Declaration of Helsinki (2008) of the Word Medical Association. Written informed consent was obtained from all participants. The study was approved by the Ethics and Clinical Investigation Committee of the Regional University Hospital of Malaga. All experimental mouse procedures were approved by either the Institutional Animal Care Committee of the Andalusian Center of Molecular Biology and Regenerative Medicine (CABIMER) or by the ethic committee of the University of Malaga, Biomedical Research Institute of Málaga (IBIMA) and performed according to the Spanish law on animal use RD 53/2013.

### Clinical Samples

Blood samples were selected from a cohort of the Pizarra Study [32]. Briefly, the inclusion age was 18 to 65 years and individuals were excluded from the study if they were institutionalized for any reason, pregnant, or had a severe clinical or psychological disorder. Venous blood samples were collected in fasting conditions and stored in a biobank at the Regional University Hospital of Malaga (Spain). Subcutaneous adipose tissue biopsies were obtained from patients with morbid obesity who underwent bariatric surgery and from non-obese patients underwent to laparoscopic surgery for hiatus hernia at the Regional University Hospital of Malaga. Age and gender of patients in both samplings were not significantly different from the population distribution. Clinical characteristics are provided in **Table 1** and **Table 2**. The research was carried out in accordance with the Declaration of Helsinki (2008) of the Word Medical Association. Written informed consent was obtained from all participants. The studies were approved by the Ethics and Clinical Investigation Committee of the Regional University Hospital of Malaga.

**Table 1.**
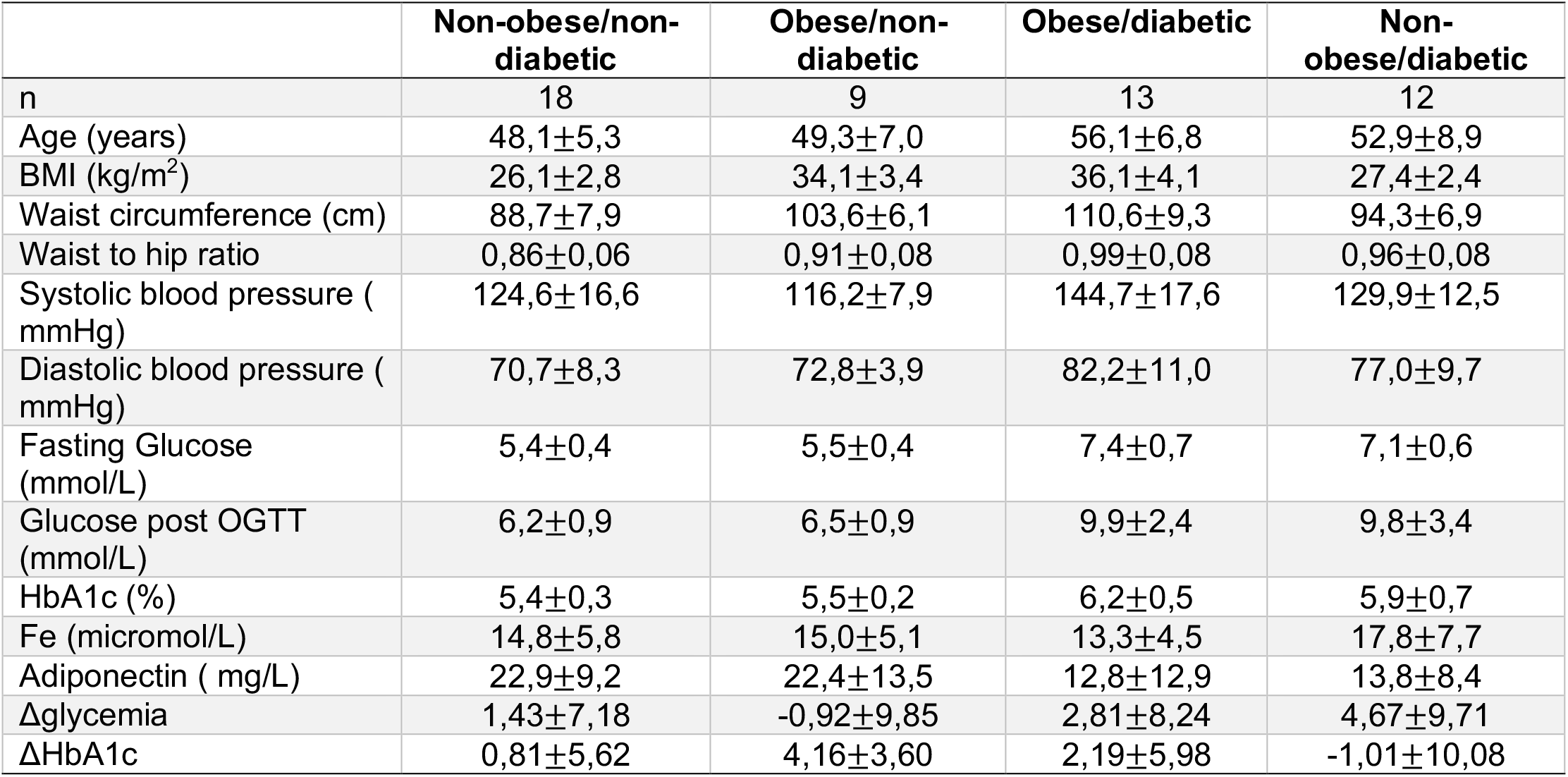
Clinical characteristic of Pizzara subjects who donated blood samples

**Table 2.** Clinical characteristic of subjects whom underwent an adipose tissue biopsy

|  | **Non-obese/non-diabetic** | **Obese/non-diabetic** | **Obese/diabetic** | **Non-obese/diabetic** |
| --- | --- | --- | --- | --- |
| n | 8 | 8 | 8 | 3 |
| Age (years) | 47.7±9.2 | 40.8±8.6 | 49.0±6.9 | 51.3±8.9 |
| BMI (Kg/m<sup>2</sup>) | 26.7±2.8 | 53.4±7.7 | 53.9±8.6 | 25.3±3.5 |
| Waist (cm) | 94.4±5.2 | 139.8±20.0 | 144.2±14.0 | 107.3±2.5 |
| Waist to Hip ratio | 0.93±0.02 | 0.91±0.06 | 0.94±0.11 | 1.12±0.08 |
| Systolic blood pressure (mmHg) | 123.0±10.4 | 120.3±15.3 | 144.6±21.5 | 127.5±28.9 |
| Diastolic blood pressure (mmHg) | 80.6±6.8 | 84.0±5.2 | 88.6±10.2 | 71.5±4.9 |
| Fasting Glucose (mmol/L) | 4.86±0.64 | 5.25±0.42 | 7.78±3.23 | 9.17±2.86 |
| HbA1c (%) | 5.6±0.4 | 5.5±0.2 | 7.2±1.3 | 7.5±1.2 |
| Fe (µmol/L) | 17.5±9.8 | 11.9±3.7 | 10.9±4.5 | 13.8±2.3 |
| Adiponectin (mg/dl) | 15.2±6.5 | 18.9±8.2 | 10.8±8.2 | 8.4±0.9 |

### Mice

All experimental mouse procedures were approved by either the Institutional Animal Care Committee of the Andalusian Center of Molecular Biology and Regenerative Medicine (CABIMER) or by the ethic committee of the University of Malaga, Biomedical Research Institute of Málaga (IBIMA) and performed according to the Spanish law on animal use RD 53/2013. Animal studies were performed in compliance with the ARRIVE guidelines [33]. C57BL/6J mice were purchased either from Charles Rivers (L’Arbresle Cedex, France) or Janvier Labs (Saint-Berthevin Cedex, France). Mice were housed in ventilated plastic cages and maintained on a 12-h light-dark cycle with ad libitum access to pellet chow and water. For diet-induced obesity studies, 8 weeks-old male mice were housed in individual cages. Two groups of fifteen mice were exposed to high fat diet (D12451 Research Diets Inc, New Brunswick, NJ, USA), containing 45% of Kcal from saturated fat, for 15 weeks. In parallel, two groups of fifteen mice were fed a control diet containing 10% of Kcal of fat (D12450 Research Diet Inc). Circulating glucose levels were measured from tail vein blood samples using an Optium Xceed glucometer (Abbott Cientifica SA, Barcelona, Spain). Insulin tolerance tests (ITT) and glucose tolerance test (OGTT) were performed as previously described [34]. In parallel, some mice were also used for organ extractions that included the brain, white adipose tissue (WAT) as well as pancreatic islets and processed for immunofluorescence analysis or RNA extraction.

### Primary hypothalamic astrocytes culture

Primary hypothalamic astrocytes were isolated and cultured as previously described [35, 36]. Briefly, hypothalamus from 2 to 4-days-old C57BL/6J male mice were extracted under sterile conditions and triturated in Dulbecco’s modified Eagle’s Medium (DMEM) F-12 (Gibco, Thermo Fisher Scientific, MA, USA) containing 1% penicillin-streptomycin (Gibco). The suspension was centrifuged and the pellet resuspended in DMEM F-12 + 10% FBS + 1% antibiotics. Cells were grown in this culture media in T75 culture flasks at 37°C and 5% CO_2_. The culture medium was refreshed every 2 days. At confluence, microglia and oligodendroglia were detached by shaking flasks at 220 rpm for 16 hours at 37°C. The remaining attached astrocytes were then used for experiments.

### Primary spinal cord motoneurons isolation and culture conditions

Motor neuron cultures were prepared from embryonic 12.5-day spinal cords. Embryos were obtained from Friend leukemia virus B (FVB) mice (University of Seville Animal Core Facility). Dissection of the spinal cord from embryonic (E12.5) were followed by the separation of neurons from the spinal cord tissue through mechanical and enzymatic cleavage as described [37, 38] with modifications. Briefly, spinal cords were dissected and meninges removed. Isolated spinal cords were transferred to ice cold Leibowitz’s 15 medium (Thermo Fisher Scientifics) and fragmented. Spinal cord fragments were washed with HEPES buffer solution (137 mM NaCl, 2 mM KCl, 25 mM glucose, 25 mM HEPES buffer, pH 7.4, and 20 IU/mL penicillin plus 20 mg/mL streptomycin) and digested with 0.025% trypsin-EDTA (Gibco) for 8 min at 37°C. After digestion, spinal cords were subjected to mechanical disaggregation by shaking in Leibowitz’s L15 media containing 0.1 mg/mL DNase (Sigma-Aldrich). Additional disaggregation was carried out by careful pipetting the samples in order to obtain a dispersion at single-cell level. Cells were purified after a 4% BSA gradient (Thermo Fisher Scientifics) by centrifugation at 180 *x*g for 5 min and resuspended in Leibowitz’s 15 media. Next, cells were carefully layer over NycoPrep 1.077 gradient (Axis-shield) diluted with 50% of GHEBS buffer (137 mm NaCl, 2.7 mm KCl, 22.2 mmglucose, 25 mm HEPES buffer, pH 7.4, 20 IU/ml penicillin, 20 μg/ml streptomycin) and centrifugated at 520 *x*g for 10 min. Collected motor neurons were cultured in Neurobasal medium (Thermo Fisher Scientifics) supplemented with 2% (w/v) B27 complement, 2% (v/v) heat inactive horse serum, 125 nM L-glutamine, 50 μM ß-mercaptoethanol, 2 μM cytarabine and 10 ng/mL of CNTF, GDNF, BDNF and HGF. All factors were purchased from Peprotech (London, UK). The motor neuron culture media was changed every 48 hours, experimental conditions were carried out at day 5 of culture. To this end, media was complemented with increasing amounts of condition media obtained from astrocyte cultures silenced or not for HMG20A. In order to evaluate survival and measure cytotoxicity, a high throughput screening was carried out by using an *Imagexpress micro system* and *MetaXpress* equipment (MDS Analytical Technologies) with a LIVE/DEAD Viability/Cytotoxicity assay for animal cells (Molecular Probes/ThermoFisher Scientific, Madrid, Spain). The number of dead cells were counted in 4 independent experiments.

### Astrocyte *in vitro* treatment

Primary hypothalamic astrocytes were cultured in the presence of: 1) low (3 mM), normal (17.5 mM) or high (30 mM) glucose concentrations for 1, 24 and 48 hours, 2) 0.5 mM palmitate for up to 48 h or 3) 100 nM ORY-1001 (Deltaclon, Madrid, Spain) for 24, 48 and 72 hours. These experiments were design as such that all samples were collected at the same final time point, including the control. Cells were then processed for either RNA extraction or immunofluorescence analysis.

### RNA interference

Primary astrocytes were transfected with either 50 μmol of a *HMG20A* small interfering (si)RNA or a luciferase siRNA (Sigma-Aldrich) as previously described [17]. Cells were seeded at a density of either 2 x 10^5^ cells per well in 6-well plates or 2 x 10^4^ cells per well on glass cover slips in 24-well plates for RNA extraction or for immunofluorescence, respectively. Twenty-four hours later cells were transfected with the siRNAs using Oligofectamine (Life Technologies) according to the manufacturer’s instructions. Seventy-two hours after transfection, media was collected for either measuring lactate release or for use as conditioned media while cells were processed for RNA isolation.

### RNA extraction and quantitative real-time PCR

Total RNA was extracted and purified from blood samples using the PAXgene Blood RNA Kit (PreAnalytiX GmbH, Hombrechtikon, CH) according to manufacturer’s instructions. Qiagen RNeasy Mini and Micro kits (Qiagen, Madrid, Spain) were used for the extraction of total RNA from the brain, hypothalamus, white adipose tissue, pancreatic islets and astrocytes. Complementary DNA using 0.5 to 1 μg RNA was synthesized using the Superscript III Reverse Transcriptase (Invitrogen-Thermo Fisher Scientific, Madrid, Spain). The RT-PCR was performed on individual cDNAs using SYBR green (Roche) [39]. Gene-specific primers were designed using Primer3Web (https://primer3.ut.ee/) and the sequences are listed in **Table S1**. Expression levels were normalized to various housekeeping genes including *β-ACTIN, HISTONE, Cyclophilin* and *Gapdh.* The relative gene expression was calculated using the 2^-ΔΔCt^ method [17].

### RNA-Sequencing

Quality of total RNA extracted from primary hypothalamic astrocytes treated or not with siHMG20A was verified using nanoRNA kit on the Agilent 2100 bioanalyzer. Quantification of RNA was evaluated by Qubit (Qubit™ RNA HS Assay). Purified RNA (150 ng) with RNA integrity number higher than 7 was used for library preparation using the TruSeq Stranded mRNA Library Preparation Kit (Illumina, San Diego, CA, USA) and sequenced on a NextSEq500 (Illumina). Two biological replicates per condition were sequenced. A mean of 30 million paired-end reads of 75 bp were generated for each sample.

RNA-seq data were initially filtered using the FASTQ Toolkit (v1.0.0) program. RNA-seq reads were aligned to the mouse reference genome mm9 using subjunc function from Rsubread package (v1.28.1) with TH1=2 and unique=TRUE parameters. The downstream analysis was performed on bamfiles with duplicates removed using the samtools (v0.1.19) rmdup command. FeatureCounts() function from Rsubread package was used to assign reads to UCSC mm9 KnownGenes (miRNAs were discarded from the analysis) using GTF.featureType=“exon”, GTF.attrType=“gene_id”, isPairedEnd = TRUE and strandSpecific = 2 parameters on duplicate removed bamfiles. Then differential gene expression analysis was performed using the voom/limma (v.3.34.9) and edgeR (v.3.20.9) Bioconductor packages. Genes that were expressed at ≥ 1 counts per million counted reads in all samples were analyzed, resulting in 12632 genes out of 21761 UCSC KnownGene genes. CalcNormFactors() function using TMM method was used to normalize samples. An adjusted *p*-value < 0.05 was determined as threshold for multiple comparisons.

Pathway analysis was performed using Gene Set Enrichment Analysis (GSEA). First LCPM (log2(CPM)) was calculated for all 12632 expressed genes for each replicate. Z-Score matrix was created and GSEA (v.2.0) analysis was performed with 1000 permutations. Normalized enrichment score (NES) and false discovery rate (FDR) were used to quantify enrichment magnitude and statistical significance, respectively. Protein interaction analysis using annotated protein-coding genes was performed using STRING [40].

### Lactate concentrations in the culture media

Lactate levels were measured in the media of primary hypothalamic astrocyte cultures 72 hours after siCT or siHMG20A transfection. One ml of medium was removed from each culture and lyophilized. Secreted lactate levels were determined using a Lactate assay kit following the manufacturer’s instructions (EnzyChrom, BioAssay Systems, Hayward, CA, USA).

### Cell death and mitochondrial metabolism

Cell death (apoptosis) was measured using the Cell Death Detection ELISA kit (Roche Diagnostics, Madrid, Spain) while mitochondrial metabolism was assessed using the MTT assay, according to the manufacturer’s recommendations (Roche, Spain). A Seahorse Cell Mito Stress Test recorded using a XF24 Extracellular Flux Analyzer (Agilent) was performed, as previously described to measure mitochondrial bioenergetics in primary astrocytes silenced or not with siHMG20A (5000 cells/well) [41]. At the end of the measurement, cell protein content was determined to normalize the data.

### HMG20A ELISA

Human serum levels of HMG20A were measured by ELISA, following the manufacturer’s instructions (MyBioSource, San Diego, CA, USA). The sensitivity of the method was 1 ng/ml. Absorbance in each well was measured by using a Varioskan Flash Spectral Scanning Multimode (ThermoFisher Scientific) and serum HMG20A concentrations were calculated from the standard curve. Samples were run in duplicates.

### Immunofluorescence studies

For brain immunostaining, mice were transcardially perfused with 4% PFA in 0.1M PB and, after overnight post fixation in the same fixative, brains were coronally sectioned into six series of 50 μm slices using a vibratome (Leyca). Free floating brain sections and fixed astrocyte cultures were immunostained with the following antibodies (**Table S2**): Mouse anti-NeuN (1:100, Merck Millipore, Darmstadt, Germany), Rabbit anti-GFAP (1:200, Dako, Barcelona, Spain), mouse anti-GFAP and rabbit anti-HMG20A (1:1000, Sigma-Aldrich, Madrid, Spain) or a combination of these antibodies. Secondary antibodies all used at a 1:400 dilutions were: Rhodamine Red Anti-rabbit (Jackson ImmunoResearch, Suffolk, UK), Rhodamine Red Anti-mouse (Jackson ImmunoResearch), Fluorescein Anti-rabbit (Jackson ImmunoResearch) and Flouresecein Anti-mouse (Jackson ImmunoResearch), Alexa Fluor 488 anti-mouse and Alexa Fluor 647 anti-rabbit (ThermoFisher Scientific). As negative controls primary antibodies were omitted from the reaction in the presence of all the reagents and secondary antibodies. All the negative controls were run in parallel for each immunodetection assay and always resulted in lack of signal (data not shown). Nuclei were stained with 0.0001% of 4’,6-diamidino-2-phenylindole (DAPI, Sigma-Aldrich). Epifluorescence microscopy images were acquired using either a Leica DM6000B or an Olympus X71 fluorescence microscope.

### Morphometry

To assess ORY-1001-mediated astrogliosis, images of GFAP immunostained astrocytes treated or not with the drug were acquired from 5 independent areas of each slides using a 10X objective. Each experimental conditions was performed in triplicate in 3 independent astrocyte primary cultures. Images were then analyzed using the ImageJ software by delineating edges of GFAP^+^ cells and calculating the surface area (μm^2^) of at least *46-56* images from which the area of at least 40 GFAP^+^ cells were measured. GFAP signal intensity was assessed in *337-396* cells.

### Western Blotting

Primary astrocyte cultures were processed and protein concentration determined as previously reported [36]. Western blotting was performed as previously described [41]. Antibodies and dilutions employed are provided in **Table S2**.

### Statistical Analysis

Measurements of the data was made in at least 4 independent samples. Results are presented as means ± SEM. Statistical comparisons were performed by Student’s *t*-test and all *p* values less than or equal to 0.05 were considered statistically significant. Statistical analyses were performed using either the SPSS software version 18.0 (IBM, NY, USA; clinical study) or the GraphPad Prism software version 8 (GraphPad Software, La Jolla, USA).

### Data and Resource Availability

All data generated and/or analyzed in the context of the current study are included in the published article (and its Supplementary Material), The RNA-seq raw data that support the findings of this study have been deposited in the Gene Expression Omnibus repository with accession number GSE161334.

## Results

### HMG20A is expressed in hypothalamic astrocytes

Transcript levels for *Hmg20a* were previously shown by *in situ* hybridization to be highly abundant in the outer cortex of the developing mouse brain [26] and postnatally in the dentate gyrus and pyramidal layer of the hippocampal formation (https://mouse.brain-map.org/experiment/ivt?id=357093&popup=true) [42]. Notwithstanding, the expression of the protein and cell-type distribution within the mouse adult brain has yet to be mapped. To address this point, immunofluorescence studies using a validated anti-HMG20A polyclonal antibody [17] were performed on various region of the mouse brain, focusing on its expression in astrocytes and neurons. HMG20A was detected in various area of the brain, mainly co-localizing with a subpopulation of NEUN^+^ neurons (**Figure S1**). In addition, HMG20A also colocalized with GFAP^+^ astrocytes (**Figure S2**) including the hypothalamic region with preponderance within the arcuate and ventromedial nuclei (**Figure 1A and B**). This expression pattern was confirmed at the transcript level in isolated hypothalamic astrocytes (**Figure 1C**). Interestingly, *Hmg20a* expression in these astrocytes was comparable to that of whole brain, islets and white adipose tissue (WAT) [17]. As the mean ratio of astrocytes, that comprise 20% of the total glial cell population, to neurons was recently estimated to be approximately 0.18 in the brain [43, 44], our data suggest that HMG20A is highly expressed in hypothalamic astrocytes while neurons likely express lower levels throughout the brain.

**Figure 1.**
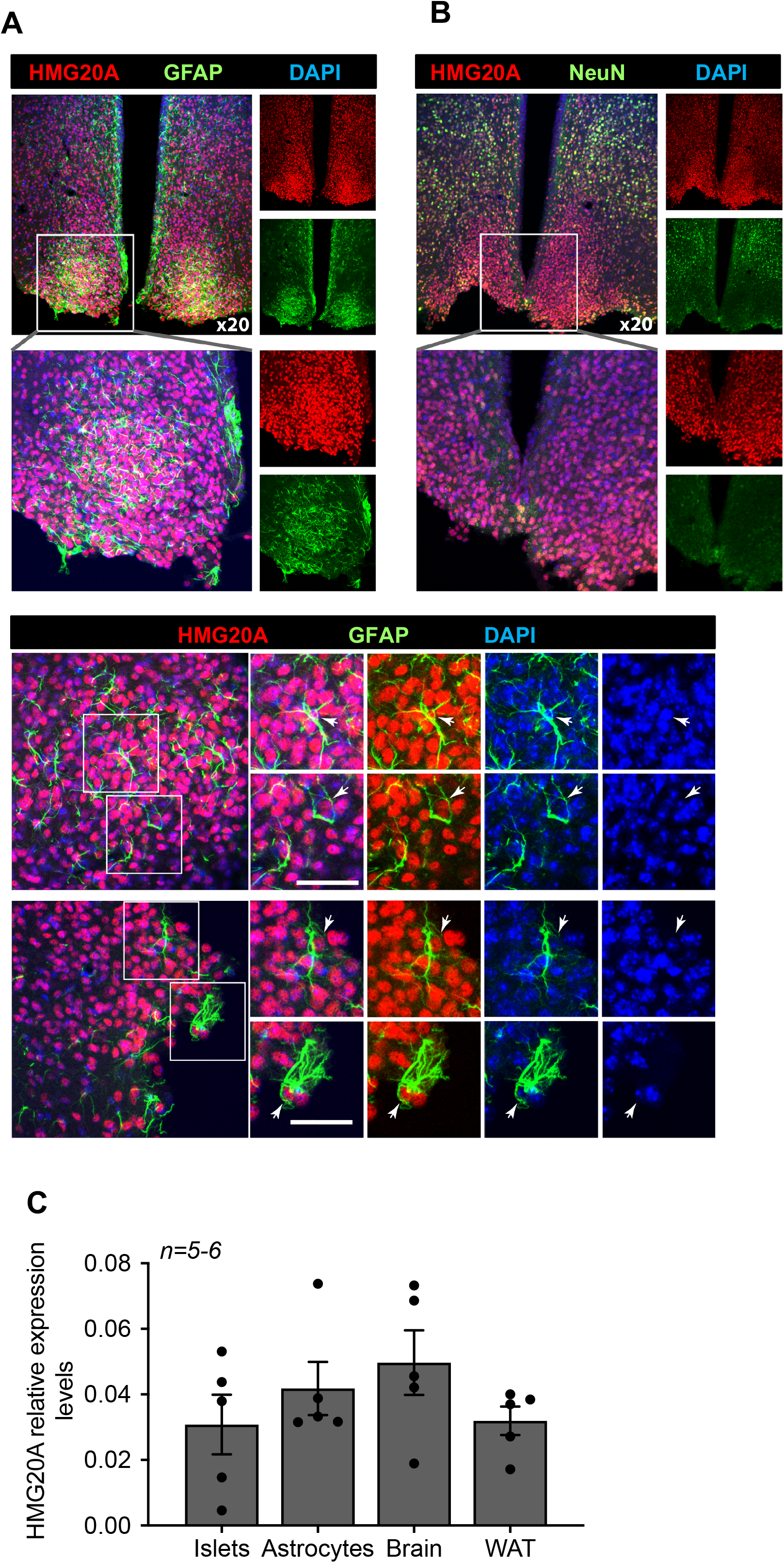
HMG20A is expressed in hypothalamic astrocytes. (**A**) Representative confocal microscopy images of free floating mouse brain sections immunostained for HMG20A (red) and GFAP, an astrocyte marker (green, upper panel left) or (**B**) NeuN a neuronal marker (green, upper panel right). Nuclei were stained using DAPI (blue). The bottom panel in (**A**) provides higher magnification images for the detection of HMG20A (red) and GFAP (green) co-expression in hypothalamic astrocytes. Arrows point to astrocytes expressing HMG20A. White bar corresponds to 50 μm. (**C**) Relative expression levels of *Hmg20a* in mouse islets, astrocytes, brain and white adipose tissue (WAT). *Hmg20a* transcript levels were normalized to those of the housekeeping gene *Gapdh* and/or *Cyclophilin. n=5-6* biological replicates analyzed in duplicate.

### Metabesity induces Hmg20a gene expression in the mouse hypothalamus as well as in human adipose tissue

We previously demonstrated that *Hmg20a* expression is enhanced in pancreatic islet beta-cells in response to increased insulin demands such as during pregnancy or pre-diabetic conditions correlating with sustained normoglycemia [17]. The hypothalamus is also a nexus in the control of glucose homeostasis via the arcuate nucleus that, in part, fine tunes insulin secretion from beta-cells while the ventromedial nucleus favours the counter regulatory response through potentiating glucagon release from alpha-cells [45]. As *Hmg20a* was found to be highly expressed in hypothalamic astrocytes, we reasoned that it may play a similar role to that observed in beta-cells and facilitate the neuronal response and/or survival to metabesity conditions. To this end, we assessed whether *Hmg20a* expression levels were altered in the hypothalamus of a high fat diet (HFD) induced-obesity and pre-diabetic mouse model that we recently developed [34]. Expression levels of *Hmg20a* along with those of the astrocytic markers *Gfap* (glial fibrillary acidic protein) and *Vimentin* were significantly increased in the hypothalamus of HFD fed mice as compared to those fed a normal chow diet (**Figure 2A**). Consistent, with obesity-induced neuroinflammation, expression of *Il1b* was higher in HFD treated mice as compared to normal chow fed mice (**Figure 2B**). Hyperglycemia and hyperlipidemia are hallmarks of metabesity [3]. As such, we sought to determine the direct contribution of either high glucose or palmitate on expression levels of *Hmg20a* in isolated hypothalamic astrocytes. High glucose transiently decreased *Hmg20a* expression at 24 hours post treatment whereas palmitate did not prompt changes in expression of the factor (**Figure 2C and D**) suggesting that glucose but not lipids may modulate expression of *Hmg20a.*

**Figure 2.**
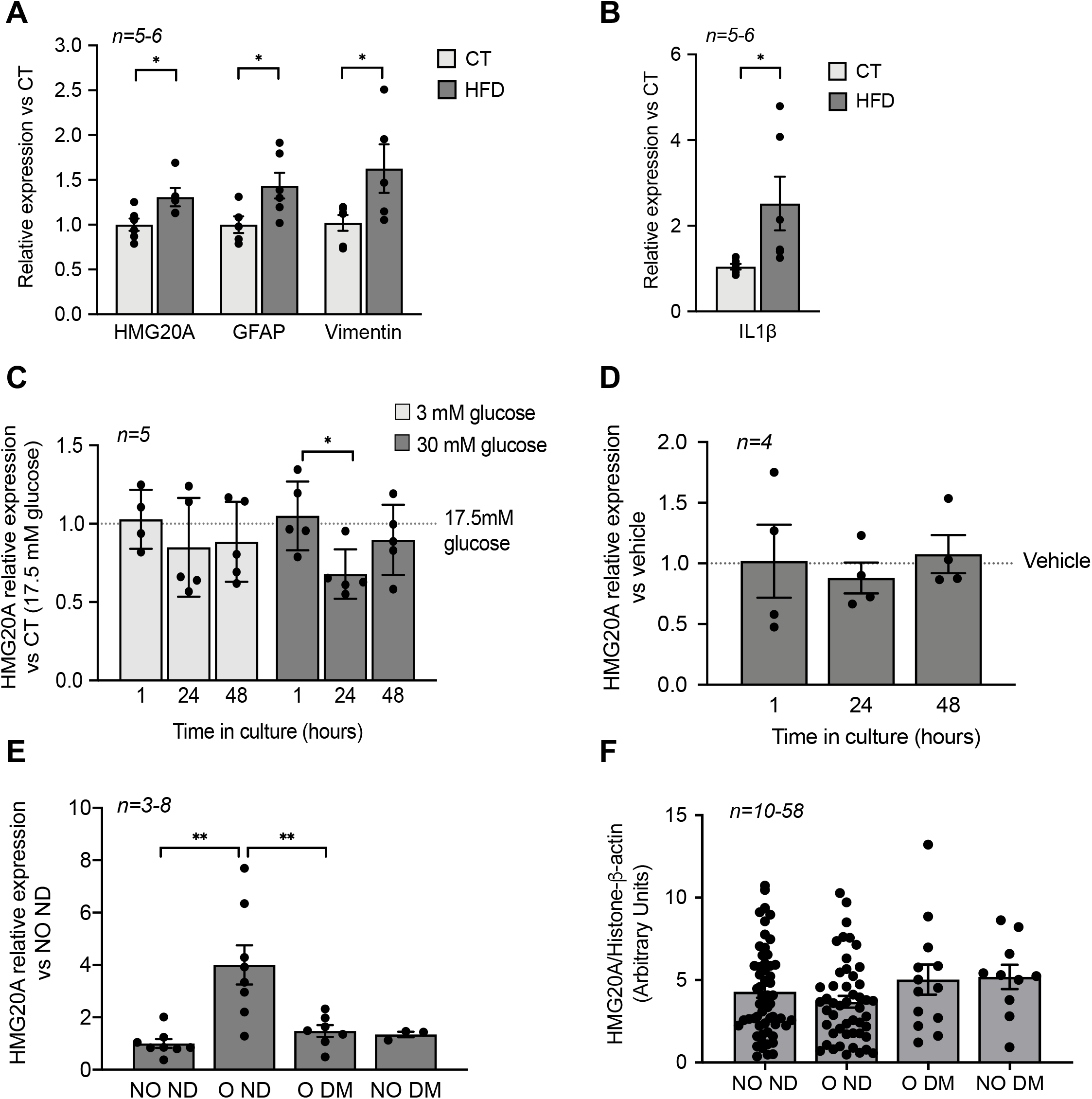
HMG20A levels are increased by physiological/metabolic stresses in both mouse and human. (**A**) Relative expression levels of *hmg20a, Gfap, Vimentin* and (**B**) *Il1b* in hypothalamus extracted from mice fed either a normal diet (CT) or a high fat diet (HFD). Transcript levels were normalized to those of the housekeeping gene b-Actin. Values are referred to the average expression levels detected in CT mice. *n=5-6* biological replicates analyzed in duplicate. *p<0.05, unpaired *t*-test CT versus HFD. Relative expression levels of *Hmg20a* in isolated mouse astrocytes cultured in either (**C**) increasing concentration of glucose or (**D**) 0.5 mM palmitate for 1, 24 and 48 hours. Transcript levels were normalized to those of the housekeeping gene *Gapdh* and/or *Cyclophilin.* Values are referred to the average expression levels detected in astrocytes cultured either in 17.5 mM glucose (**C**) or untreated (vehicle, **D**). *n=4-5* biological replicates analyzed in duplicate. *p<0.05, unpaired *t*-test 1 versus 24 hours. (**E**) *HMG20A* expression levels were measured in subcutaneous white adipose tissue biopsies that were obtained from non-obese/non-diabetic (NO ND), obese/non-diabetic (O ND), obese/diabetic (O DM) and non-obese/diabetic (NO DM). The expression of *HMG20A* was corrected to those of the housekeeping gene *HISTONE.* Values are referred to the average expression detected NO ND individuals. *n=3-8* individuals per group, analyzed in duplicate. ** p<0.01, unpaired *t-*test NO ND versus O ND or O DM. (**F**) Leukocytes were isolated from blood samples extracted from non-obese/non-diabetic (NO ND), obese/non-diabetic (O ND), obese/diabetic (O DM) and non-obese/diabetic (NO DM) and relative expression levels of *HMG20A* were assessed by quantitative PCR. The expression of HMG20A was corrected to those of the housekeeping genes *HISTONE* and *B-ACTIN. n=10-58* individuals per group analyzed in duplicate.

We previously demonstrated that *HMG20A* expression levels are blunted in islets isolated from human T2DM donors [17]. We therefore contemplated whether *HMG20A* expression is increased as a tissue adaptive response to glucose in order to enhance cell function, as assessed by higher levels in brain of prediabetic mice or islet beta-cells of gestating female but collapses in chronic pathophysiological conditions such as DM, levels resulting in cellular disarray. To examine this possibility in humans, we assessed the expression levels of *HMG20A* in subcutaneous adipose tissue biopsies taken from donors who were: 1) non-obese/non diabetic (NO ND), 2) obese/non-diabetic (O ND), 3) obese/diabetic (O DM) and 4) non-obese/diabetic (NO DM) (**Table 2**). We elected for WAT as *HMG20A* is expressed in this glucose-homeostasis regulatory tissue (**Figure 1C**) for which biopsies can be easily acquired as compared to the brain. Consistent with our premise, *HMG20A* expression levels were significantly higher in WAT of obese/non-diabetic as compared to either non-obese non diabetic or diabetic individuals independent of weight (**Figure 2E**). In order to extend these findings as well as to identify a possible blood diagnostic marker for DM, HMG20A levels were also evaluated in blood serum and leukocytes of healthy or diabetic donors (**Table 1**). No significant differences in protein or transcript levels were detected in serum or leukocytes among the various groups (**Figure S3 and Figure 2F**). In summary, HMG20A expression is modulated in mouse hypthalamus and human WAT in response to metabesity conditions suggesting a potential adaptive mechanism similar to pancreatic islets [17].

### HMG20A regulates pathways involved in astrogliosis

In order to decipher the regulatory function of HMG20A in astrocytes, the chromatin factor was silenced and RNA-seq was carried out to identify differentially expressed genes (DEGs) as well as their downstream functional consequences. A 60% repression was achieved using a previously validated siHMG20A when compared to a control luciferase siRNA (siCT) (**Figure 3A**) [17]. Transcriptome analysis revealed that 91 transcripts were upregulated and 245 were downregulated in siHMG20A transfected astrocytes as compared to siCT astrocytes (**Figure 3B**). Integrated bioinformatic tools using different algorithms were applied to analyse DEGs in order to reveal key pathways altered by HMG20A repression. Several gene ontology (GO) biological process (BP) related to cell migration and adhesion as well as to fatty acid and sterol biosynthetic processes were down-regulated subsequent to HMG20A depletion in astrocytes (**Figure 3C**). Interestingly, most DEGs within these BPs were down-regulated as exemplified by genes implicated in the steroid biosynthesis pathway (**Figure S4**). This premise was reinforced using an unbiased STRING analysis that revealed 4 main biological clusters of down-regulated genes (stress response and inflammation, growth factors and signal transductions, cytoskeleton-integrin connection and lipid metabolism) while none were found for up-regulated genes (**Figure S5**). The GSEA algorithm was next applied in order to gain further insight into the functional clustering of DEGs. In line with both GO and STRING, GSEA revealed common functional groups including inflammatory response, cholesterol homeostasis and epithelial to mesenchymal transition (EMT) (**Figure 3D-F**) as well as several cytokine related pathways such as TGFB1 and IL2 (**Figure S6**) that were down regulated by siHMG20A. Of interest, these pathways and associated DEGs (heatmaps, **Figure 3D-F**) have been linked to astrogliosis and neuron viability. For example, TGFB1 was shown to be both a para- and auto-crine signalling molecule that stimulate the reactive astrocyte phenotypic state while VCAM1 and IGFBP3 facilitate EMT, a cellular process important for transitioning between non-reactive and reactivated astrocytes [15, 46, 47]. Similarly, astrocytic expression of SREBF2 the central transcription factor coordinating the expression of cholesterol biosynthesis genes as well as SLC1A2 a glutamate transporter were shown to be essential for neuronal function an viability [48-50]. Repression of these genes in siHMG20A treated astrocytes was confirmed by QT-PCR (**Figure 3G**). Taken together these results indicate that HMG20A coordinates key genetic components governing astrocyte phenotypic state.

**Figure 3.**
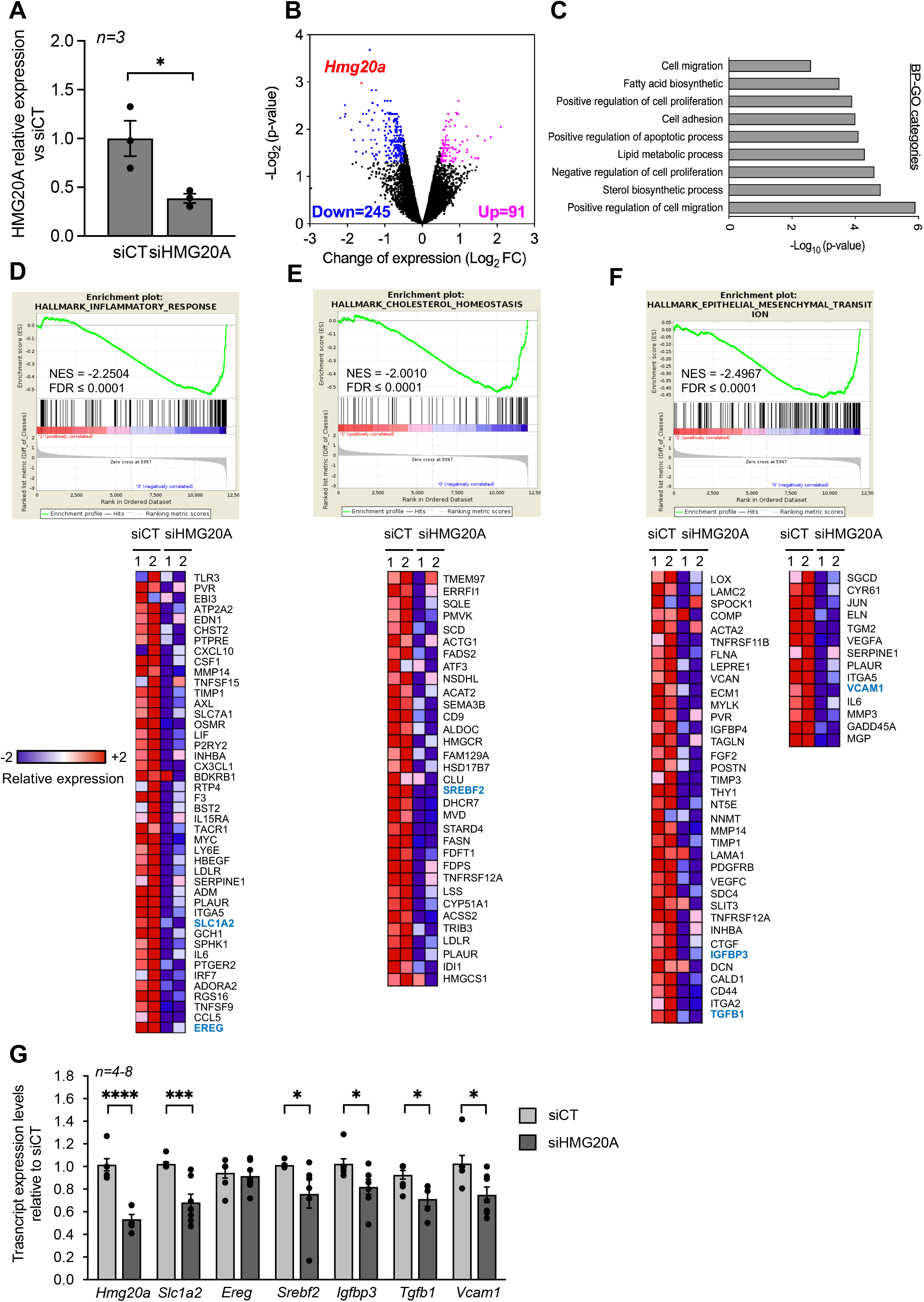
HMG20A silencing represses signaling pathways linked to astrogliosis. (**A**) Relative expression levels of *Hmg20a* in astrocytes exposed to either a control luciferase siRNA (siCT) or a siRNA targeted to HMG20A (siHMG20A). Transcript levels were normalized to those of the housekeeping gene *Gapdh* and/or *Cyclophilin.* Values are referred to the average expression levels detected in siCT. *n=3* biological replicates of primary astrocyte preparations pooled from 3-6 mice and analyzed in duplicate. *p<0.05, unpaired t-test siCT versus siHMG20A. (**B**) Volcano plot representation of differentially regulated genes after HMG20A silencing, analyzed by RNA-seq. Genes significantly (p-value < 0.05) downregulated are marked in blue (Log2FC ≥ - 0.5) and the upregulated ones in purple (Log2FC ≥ 0.5). *Hmg20A* is highlighted as a red dot. (**C**) Biological process (BP)-GO categories enriched after HMG20A silencing, analyzed using the David software. (**D**) (**E**) (**F**) Representative enriched pathways (upper panels) related to astrogliosis. The analysis was performed using the GSEA software. Normalized Enrichment Score (NES) and False Discovery Rate (FDR) are indicated within the plots. Lower panels depict heatmaps of the relative expression of those genes significantly downregulated after HMG20A silencing in each enriched pathways in two independent RNAseq. (**G**) Confirmatory QT-PRC of selected target genes from the various pathways altered by HMG20A repression. Transcript levels were normalized to those of the housekeeping gene *Gapdh* and/or *Cyclophilin.*

### *Hmg20a* repression in astrocytes blunts mitochondrial metabolism sensitizing cells to apoptosis

Although up-regulated genes did not cluster into functional hubs, individual genes such as *Bcl2l11 (aka Bim)* were significantly increased in siHMG20A-treated cells. *Bcl2l11* is of particular interest in view of its outer mitochondrial membrane localization and its role in regulating mitochondrial bioenergetics as well as BAX-dependent cell apoptosis [51]. As mitochondrial dysfunction, a hallmark of increased *Bcl2l11* expression, was recently found to impede astrogliosis and to enhance neuronal cell death [52], we assessed mitochondrial metabolism in siHMG20A-repressed astrocytes. We first confirmed that *Bcl2l11* expression was increased in siHMG20A knockdown astrocytes (**Figure 4A**). These cells displayed a 30% reduction in NADH oxidation as assessed by the MTT assay suggestive of lower mitochondrial bioenergetics (**Figure 4B**). To further explore this premise, we measured oxygen consumption rate (OCR) and extracellular acidification rate (ECAR) in astrocytes silenced or not for *Hmg20a* using the seahorse XF-24 bioanalyzer. Basal and maximal respiration (following the addition of FCCP) were significantly reduced while proton leakage (following the addition of oligomycin) was unchanged in siHMG20A-transfected astrocytes as compared to siCT-treated cells (**Figure 4C and D)**. The extracellular acidification rate (ECAR) was not altered consistent with sustained lactate secretion (**Figure 4E and F**). In line with lower ATP production, *Hmg20a* knockdown astrocytes were more susceptible to apoptosis (**Figure 4G**). Taken together these results indicate that HMG20A likely *via* BIM is required to sustain normal mitochondrial oxygen consumption and astrocyte survival.

**Figure 4.**
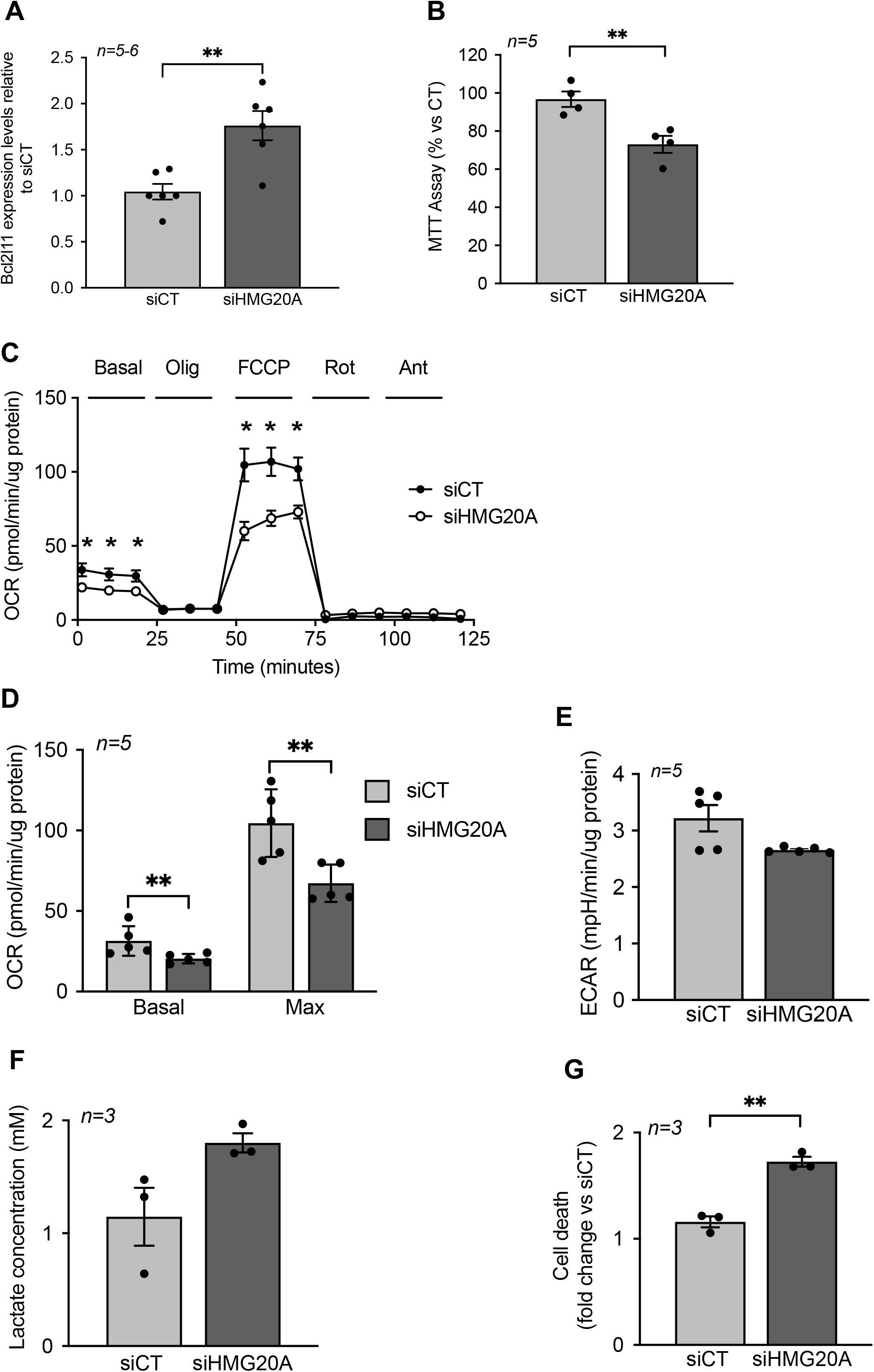
HMG20A silencing alters mitochondrial bioenergetics sensitizing astrocytes to apoptosis. (**A**) *Bcl2ll1* transcript levels in siCT or siHMG20A treated astrocytes. Expression levels of *Bcl2l11* were normalized to the housekeeping gene *Gapdh. n=5* independent experiments, ** p<0.01 unpaired Student *t-*test siCT versus siHMG20A. (**B**) Knockdown of HMG20A in astrocytes significantly decreased mitochondrial metabolism as assessed by MTT assay. Seahorse analysis showing (**C**) OCR profile plot, (**D**) Basal and maximal respiration and (**E**) ECAR in astrocytes silenced or not for HMG20A. Oligo; Oligomycin, FCCP; Carbonyl cyanide-p-trifluoromethoxyphenylhydrazone, Rot; Rotenone, Ant, Antimycin A. *n=5* (independent primary cultures obtained from 4 male pups of 2-4 days old)*, \** p<0.05 unpaired *t-*test siCT versus siHMG20A. (**F**) Lactate secretion and (**G**) Apoptosis was assessed subsequent to siRNA-mediated HMG20A silencing in astrocytes. (**B**), (**F**) and (**G**) Data are referred to the average value detected in non-silenced astrocytes (siCT). For each condition, 3-4 biological replicates (independent primary cultures obtained from 4 male pups of 2-4 days old) were analyzed in duplicate. ** p<0.01 unpaired Student *t-*test siCT versus siHMG20A.

### Hmg20a expression in astrocytes is mandatory for motoneuron survival

Our RNA-seq analysis pinpoint to a role of HMG20A in coordinating key molecular pathways transitioning astrocytes between a non-reactive and reactive phenotype, both mandatory for neuronal survival, pending environmental cues. As such, to functionally translate this molecular premise, isolated primary mouse motoneurons were cultured with increasing amounts of conditioned media derived from either siCT or siHMG20A-treated astrocytes and cell viability was assessed 24 hours post incubation (**Figure 5A**). In cultures using conditioned media from HMG20A-silenced astrocytes, we observed a media mixture-dependent increase in motoneuron apoptosis reaching a maximum of 3 fold when using a 50% ratio (**Figure 5B**) validating the essentiality of *Hmg20a* expression in astrocytes to support motoneurons survival likely via extracellular factors. Accordingly, further analysis of our RNA-seq data revealed that several GO cellular compartments with genes related to the extracellular space were significantly altered in siHMG20A-treated astrocytes (**Figure 5C**). More specifically, we found *an armada* of growth factors some of which are key in neuron survival and function such as *Gdf15, Il11 and Tgfb1* to be significantly blunted in siHMG20A-treated astrocytes [53-55] (**Figure 5D**). Oddly, *Gdf10* was the only up-regulated growth factor in HMG20A silenced astrocytes. Taken together these results confirm that the molecular alterations induced by HMG20A depletion not only impacts astrocytes phenotypic polarization but also their capacity to convey survival signals to motoneurons.

**Figure 5.**
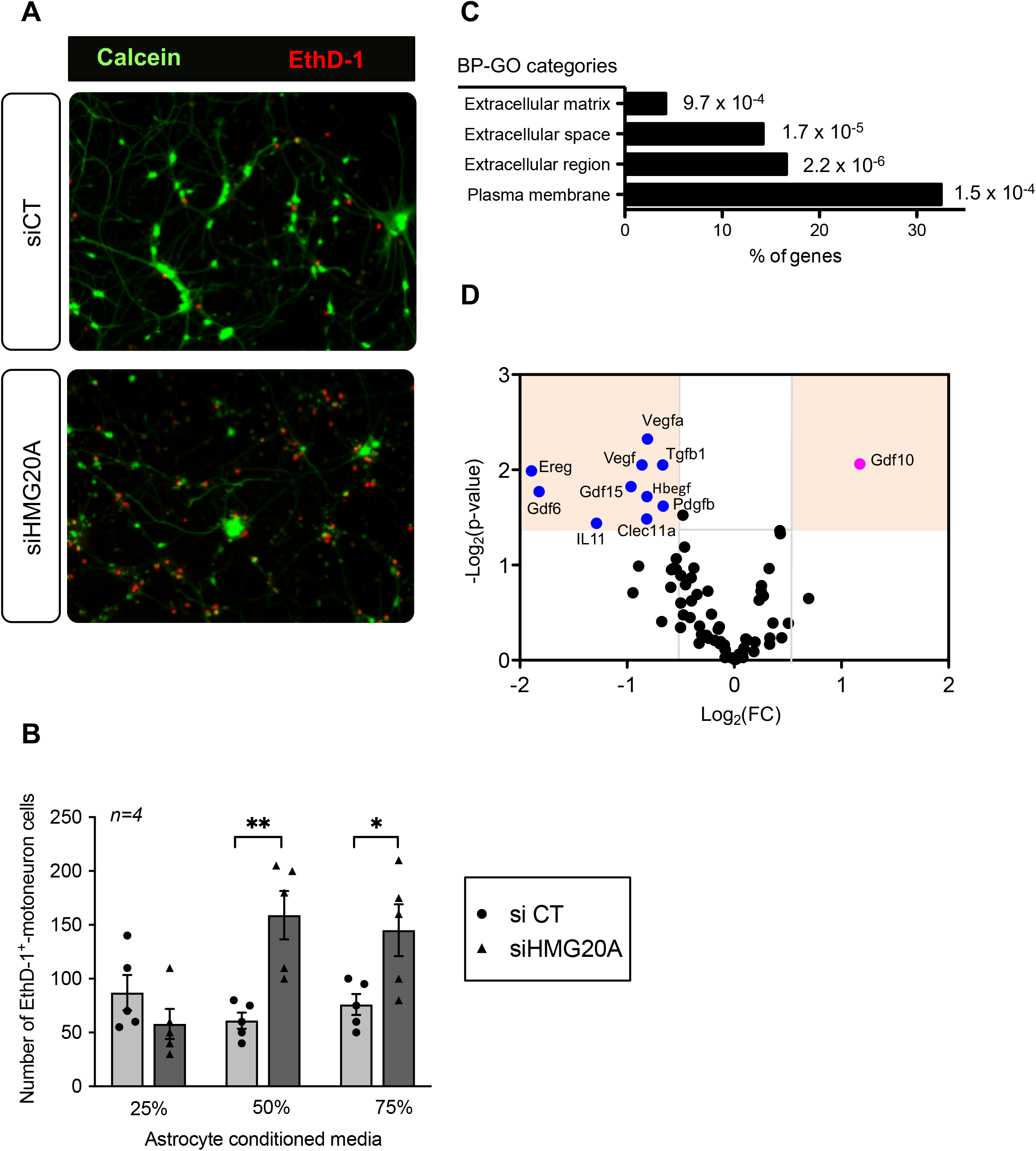
HMG20A coordinates expression of astrocyte derived extracellular factors involved in neuronal survival. (**A**) Representative immunofluorescent images of mouse spinal cord motoneurons cultured for 24 hours in increasing percentage of conditioned media derived from primary astrocytes silenced for HMG20A (siHMG20A) or not (siCT). Live cells are stained with Calcein (green) while dying cells label for Eth-D1 (red). (**B**) Quantification of motoneuron cell death after growth in the indicated % of astrocyte conditioned media; control siCT astrocytes (black circles) or HMG20A silenced siHMG20A astrocytes (Black triangles). n=6 biological replicates (independent spinal cord motoneurons primary cultures). * *p*<0.05 and ** *p*<0.01, unpaired *t-*test siCT versus siHMG20A. (**C**) GO Cellular compartment of genes differentially regulated by siHMG20A in astrocytes. % of genes of the indicated categories is represented. *p-*value of the enrichment is also provided. (**D**) Vulcano plot showing expression change (logs(FC) and *p*-value (-log10(p-value)) (siHMG20a versus siCT) of all expressed growth factors encoding genes.

### Pharmacological inhibition of the HMG20A-regulated LSD1/CoREST complex induces astrogliosis

A key molecular regulatory function of HMG20A is to counteract HMG20B-mediated activation of the LSD1/CoREST complex that facilitates demethylation of H3K4me1/2 via LSD1 (*aka* KDM1A) resulting in the repression of neuron-specific genes [26]. More recently, we demonstrated that inhibition of HMG20A blunted EMT, a process also implicating LSD1 [31]. Since astrogliosis was defined as an EMT-like process [15], we posit that pharmacologically mimicking the action of HMG20A in repressing the LSD1/CoREST complex should induce astrogliosis. Towards addressing this premise, primary astrocytes were treated with ORY-1001, a potent and selective covalent inhibitor of LSD1 [56]. ORY-1001 treated GFAP^+^-astrocytes displayed time-dependent hypertrophy with increasing cellular processes, doubling their surface area at 48-72 hours post-treatment as compared to untreated cells collected at 72 hours post-treatment (**Figure 6A and B**). Upregulation of GFAP as well as VIMENTIN has been associated with astrogliosis in multiple CNS diseases [12]. Although we did not detect any differences in levels of both proteins by Western blot analysis (**Figure S7**), quantification of GFAP immunostaining signal intensity at the single cell level revealed a significant time-dependent increase in ORY-1001 reactive astrocytes as compared to untreated cells (**Figure 6C**). As hypertrophy and extended cellular processes as well as increased GFAP expression are hallmark of astrogliosis, we conclude that HMG20A through regulation of the LSD1/CoREST complex contributes to fine tuning the phenotypic state of astrocytes to the physiological environment.

**Figure 6.**
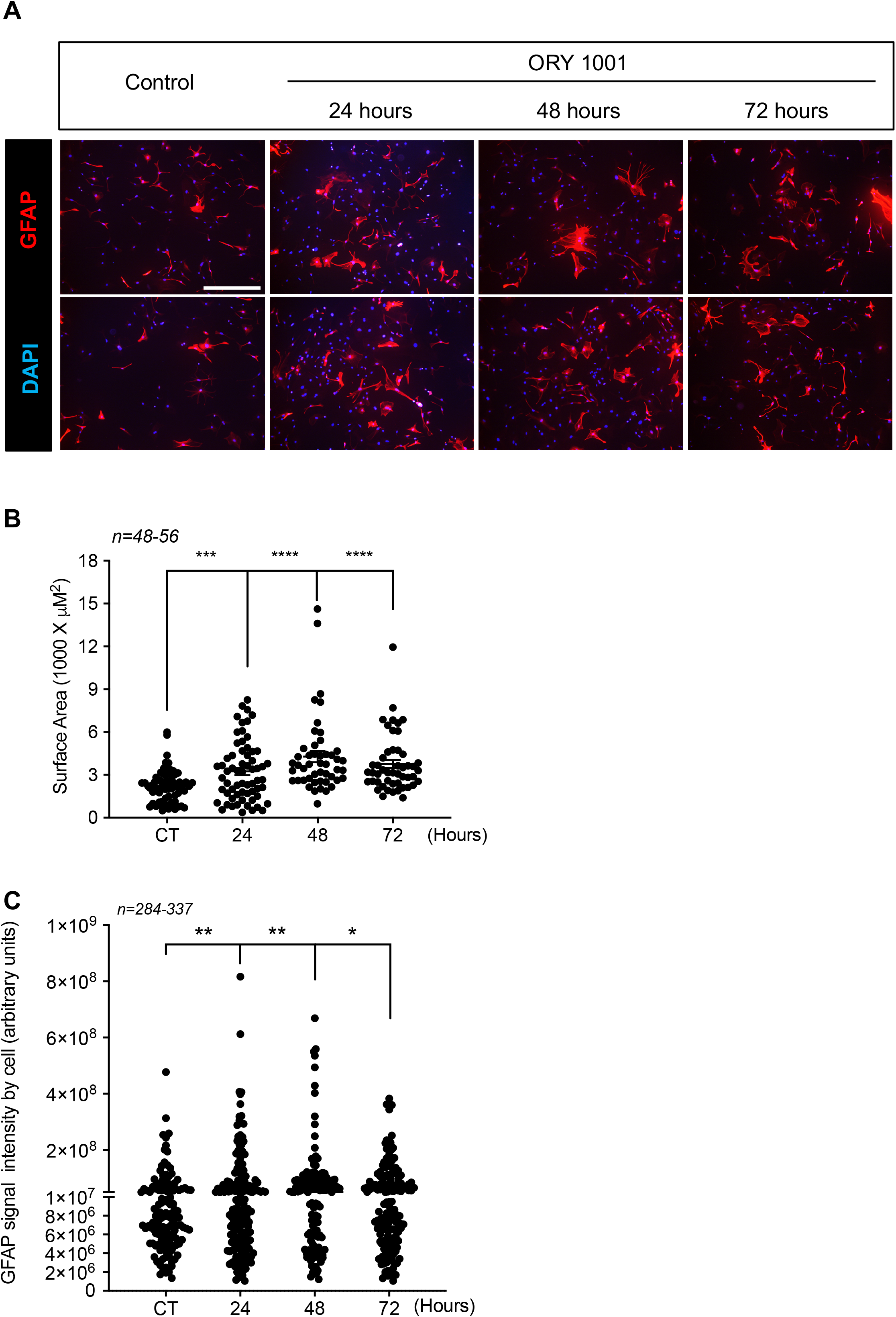
The LSD1 inhibitor ORY1001 mimics HMG20A-mediated astrogliosis. (**A**) Representative immunofluorescent images of primary astrocytes treated or not for 24, 48 and 72 hours with ORY1001. Cells were immunostained for GFAP (red) and nuclei with DAPI (blue). Two images are provided for each time point. Scale bar: 500 um. (**B**) Quantification of reactive astrocytes (i.e. astrogliosis) in (**A**) as assessed by GFAP expression and average cell surface area. *n=46-56* dot representing an image from which the area of at least 40 GFAP^+^ cells were measured from 3 independent slides/experiments. (**C**) Quantification of reactive astrocytes in (**A**) as assessed by GFAP intensity. *n=337-396* cells from 3 independent slides/experiments. **** p<0.0001, *** p<0.001, ** p<0.01 and *p<0.05 unpaired two-tailed *t-*test CT versus 48 and 72 hours.

## DISCUSSION

The overall aim of the current study was to extend our recent findings, demonstrating that the metabesity-linked chromatin factor HMG20A coordinates pancreatic islet betacell maturation and functional output in response to altered glucose levels [17], to astrocytes, the major glucose-metabolising cell subpopulation within the brain hypothalamus. We find that: (1) HMG20A is highly expressed in astrocytes within the arcuate and ventromedial nuclei of the hypothalamus cells, the nexus of brain-regulated glucose homeostasis, (2) *Hmg20a* levels are increased in hypothalamic astrocytes of pre-diabetic and obese mice, and in line with its involvement in tissue adaptive response, correlating with higher levels in WAT samples of obese non-diabetic individuals, 3) *HMG20A* levels collapse in obese diabetic patients as compared to obese non-diabetic individuals, (4) HMG20A coordinates expression of strategic hubs involved in astrocyte reactivity, (5) HMG20A depletion impairs astrocyte function and neuron survival and (6) pharmacological inhibition of the LSD1/CoREST complex, mimicking HMG20A mechanism of action, induces astrogliosis. Therefore, we conclude that HMG20A is a key molecular regulator of astrogliosis fine-tuning the reactivity degree in response to physiological cues with the ultimate purpose to protect and/or control damage of neurons.

Our results indicate that in the context of metabesity neither glucose nor lipids individually are influencers contributing to enhanced *Hmg20a* expression in astrocytes and their subsequent reactive phenotype. Although counterintuitive, in view of astrocytes intimate contact with the blood brain barrier (BBB) and function as energy sensor, our results corroborate previous studies demonstrating that under physiological conditions astrogliosis does not occur in isolation, but as part of a coordinated and multicellular response to CNS insult that includes a crosstalk with neurons and microglia [16]. As such, consumption of a HFD was shown to induce expression of IL-1beta, and IL-6 as well as TNF alpha in mouse neurons as well as to convey a pro-inflammatory phenotype to microglia within the hypothalamic arcuate nucleus in both mice and humans [7, 57, 58]. Such cytokines have been recognized as paracrine instructive signals promoting astrogliosis and thus could be mediating the HFD-induced HMG20A expression *in vivo* [46]. This is consistent with the knowledge that reactive astrocytes are found at site of CNS injury likely induced by the initial pro-inflammatory environment triggered by damaged neurons and activated microglia [12].

As discussed above, astrogliosis is an inflammatory process that initially conveys neuron protection but that long-term needs to be resorb to avoid detrimental damage. Interestingly, a negative feedback loop between the LSD1/CoREST complex and the orphan nuclear receptor NURR1 was shown to attenuate simultaneously the pro-inflammatory response in astrocytes and microglia. The authors concluded that such crosstalk blunts a feed forward amplification of the NF-κB mediated inflammatory response restoring a non-reactive phenotype to both microglia and astrocytes thereby protecting dopaminergic neurons from inflammation-induced cell death [59]. These findings are contextually relevant as a key regulatory function of HMG20A is also to counteract the effects of the LSD1/CoREST complex pinpointing to a strategic function of HMG20A in potentiating astrocyte reactivity. This premise was validated at the cellular level by demonstrating that the potent inhibitor of the LSD1/CoREST complex, ORY-1001 induced astrogliosis. At the molecular level, transcriptome profiling revealed the regulatory function of HMG20A on key genetic hubs, including the inflammatory response involved in astrogliosis. Of particular relevance for the transitory status of astrogliosis, numerous genes within the inflammatory hub that were down regulated by siHMG20A appear to encode anti-inflammatory and pro-regenerative factors (**Figure 3D**). For instance, TLR3 was shown to be induced in human astrocytes upon inflammation and when activated, to convey a neuroprotective response rather than a pro-inflammatory reaction [60]. In addition, although chronic high levels of CCL5 have been associated with MS [61], this chemokine was reported to promote an anti-inflammatory resolution-phase of macrophage reprogramming under some settings [62]. Likewise, IRF7 was shown to induce a pro-to-anti-inflammatory phenotypic shift in microglia [63]. Finally, LIF through its immunomodulatory and regenerative properties promotes survival of neurons and glia in several animal models of neurodegenerative disease, such as amyotrophic lateral sclerosis, MS, spinal cord injury and stroke [64]. These data suggest that HMG20A modulates astrogliosis, both at its initiation as well as at its reabsorption. It is therefore tempting to speculate that the differentiation/polarization status/level of astrocytes is a continuum of changes which is dictated by the interaction of HMG20A and/or NURR1 with the LSD1/CoREST complex favoring an antiinflammatory state of reactive astrocytes with pro-survival and pro-regenerative properties. Similarly, in our previous studies in islet beta cells, we detected a transitory increase in HMG20A levels during pregnancy, correlating with the adaptation phase that takes place in islets in response to increase insulin demands [17, 18]. As such, under environmental insults such as hyperglycemia, HMG20A could activate on one hand the adaptation pathways conveying maturation/functionality of beta cells in islets and and on the other hand transitory astrogliosis and neuronal protection in CNS. This molecular model could rationalize our findings that obese non-diabetic individuals have higher levels of HMG20A in WAT favoring an anti-inflammatory and pro-survival cell phenotype whereas obese diabetic patients expressed normal levels of HMG20A indicative of a non-reactive cell phenotype incapable of relaying cell protection and repair ultimately leading to neurodegeneration disease such as MS as well as other complications such as altered glucose homeostasis. This model will require further validation including the challenging task of assessing HMG20A expression levels in hypothalamus of obese nondiabetic and obese diabetic individuals.

Our molecular analysis also revealed that silencing of HMG20A repressed the EMT signaling program including the triggering factor TGF-beta. These results substantiate our previous study in which we demonstrated that HMG20A together with the LSD1/CoREST complex is required for TGF-beta-mediated EMT in mouse mammary epithelial NMuMG cells [31]. Astrogliosis was recently defined as an EMT-like process that fine tunes astrocyte reactivity in response to injury and tissue repair [15]. As such a parallel between astrogliosis and wound healing was proposed including a role of astrocytes in trans-differentiating into functional neurons, a process facilitated by VEGF [65, 66]. This growth factor along with a subset of genes implicated in cytoskeleton-integrin connection previously shown to be involved in wound healing were down regulated in siHMG20A treated astrocytes. Taken together these molecular data further support our model that HMG20A likely potentiates the degree of astrogliosis promoting an anti-inflammatory and pro-regenerative phenotype, analogous to our recent studies establishing the role of the orphan nuclear receptor LRH-1/NR5A2 in immunomodulation and trans-regeneration of islet beta cells [39, 67, 68].

We also provide functional data derived from our molecular analysis highlighting a cell autonomous role of HMG20A in promoting astrocyte viability and differentiated/reactive state. We show that HMG20A is required to sustain normal levels of mitochondrial oxygen consumption since its deficiency leads to impaired bioenergetics emphasized by reduced basal and maximum oxygen consumption, which might contribute to sensitive astrocytes to apoptosis. Of particular interest, mitochondrial dysfunction was shown to impede the generation of reactive astrocytes fostering neuronal cell death consistent with the finding that reduced reactive oxygen species (ROS) in astrocytes induced major alterations in brain energy and redox metabolism leading to impaired neuronal function and cognitive learning [52, 69]. These studies are in line with our findings that HMG20A through various cellular pathways including mitochondrial metabolism coordinates astrocyte phenotypic state. We argue that these functional consequences are likely conveyed by HMG20A-regulated BIM expression. Indeed, in addition to its known role in apoptosis, BIM was also shown to regulate mitochondrial activity. Loss of BIM was found to increase oxygen consumption characterized by higher basal and maximum respiration rates in mouse embryonic fibroblast [51] corroborating our findings that increased expression results in decreased mitochondrial bioenergetics. These results are in line with a previous study revealing that BIM potentiates mitochondrial membrane potential [70]. In parallel, the capacity of HMG20A to potentiate astrogliosis through the LSD1/CoREST complex was functionally validated using the LSD1 inhibitor ORY1001. Of high relevance, Vafidemstat (ORY2001), an oral small molecule that has been optimized for CNS indications and that acts as a covalent inhibitor of LSD1, similar to ORY1001, was shown to reduce neuroinflammation and improve cognitive behavior in animal models of AD and MS [71, 72] as well as to improve inflammation biomarkers in AD patients enrolled in a phase IIa clinical trial [73] (https://clinicaltrials.gov/ct2/show/study/NCT03867253). In view of the link between AD and T2DM [74, 75], these results, clearly suggest that targeting either HMG20A or the LSD1/CoREST complex in astrocytes may improve hypothalamus insulin-sensitivity and glucose homeostasis as well as in pancreatic islets in T2DM patients [3]. In this context, we recently provided proof-of-concept in insulin-producing cells in which short-term treatment with ORY1001 reversed the effect of HMG20A silencing on the expression of key beta cell genes involved in insulin secretion [76].

We also present functional data for a non-cell autonomous role of HMG20A in supporting motoneuron survival. Accordingly, a great number of secreted growth factors, previously shown to promote neuron survival and function [12] were down-regulated in HMG20A knockdown astrocytes. The latter suggests that astrocytes harbor a repertoire of secreted factors capable of jointly supporting neuron survival and regeneration as exemplified by the recent finding that transplanted immature astrocytes do not polarize to a pro-inflammatory reactive phenotype after CNS injury, but rather, favor neuron outgrowth and improve cell therapeutic outcomes in a Parkinson’s disease mouse model [77, 78]. Notwithstanding, a single growth factor, GDF10 *aka* bone morphogenetic protein 3 (BMP3), was significantly up-regulated while other members such as GDF6/BMP13 and GDF15/placental BMP were down-regulated in astrocytes lacking HMG20A. BMPs form a subgroup in the TGF-beta superfamily and act as powerful morphogens that play dynamic roles in the development of the brain early on in life, during which they sequentially induce neurogenesis and then astrogliogenesis [79]. In adult, both GDF6/BPM13 and GDF15/placenta BMP were found to possess potent trophic effect on motor and sensory neurons [53, 80]. Of particular importance was the finding that BPM6 conveys a protective effect against amyloid beta-induced neurotoxicity, a hallmark of AD, in rat hippocampal neurons [81] Interestingly, GDF10/BMP3 is a unique member of this family as it functions as an antagonist of the BMP pathway [82]. As such the opposing regulation of GDF6 and GDF15 as compared to GDF10 emphasises again the role of HMG20A in establishing a pro-survival and tissue repair molecular phenotype to astrocytes.

The overall data presented herein allocate a molecular and functional link between HMG20A and astrocyte reactivity. HMG20A regulates molecular hubs modulating phenotypic changes needed for astrocytes to respond to different physiological situations such as metabesity. Failure on those HMG20A-mediated processes leads to astrocyte dysfunction and ineptitude to react resulting in neuron death and to the potential development of neurodegenerative diseases such as AD as well as T2DM, for which a link has previously been established. As consequence, our study opens a venue to consider targeting HMG20A regulatory function, as exemplified by ORY1001, to treat such diseases.

## ACKNOWLEDGMENTS

We thank Ms. Eloisa Andujar Pulido and Monica Perez from the Genomic Core Facility of CABIMER as well as Dr. Maria José Quintero and Paloma Dominguez from the Cytometry and Microscopy Core Facility of CABIMER for their excellent technical and analysis support. We acknowledge the support of the pancreatic islet study group of the Spanish Association of Diabetes.

## AUTHOR CONTRIBUTIONS

P.I.L, E.F.-M, D.P., J.C.R. and B.R.G. were involved in research design. P.I.L., E.F.-M, J.M-M-G., N.C.V., V.M., E.M.V., S.Y.R.-Z., J.M.F., J.A.P.-C., S.R.C., C.C.L., E.G.F., A.M.M., M.A.D. and F.J.B.-S contributed to conducting experiments. P.I.L., E.F.-M, J.M-M-G., J.A.G.M., G.R.M., D.P., F.J.B.-S., J.C.R, and B.R.G. contributed to data analysis. A. C.-C., M.A.-A, F.M., F.J.B.-S., G.R.M. and E.G.F. supplied human samples and reagents. B.R.G. wrote the manuscript. All authors commented on the manuscript. B. R.G. is the guarantor of this work and, as such, has full access to all the data in the study and takes responsibility for the integrity of the data and the accuracy of the data analysis.

