## SUPPLEMENTAL TABLES AND FIGURES for "The Metabesity Factor HMG20A Potentiates Astrocyte Survival and Reactivity Preserving Neuronal Integrity"

Petra I. Lorenzo et al.

| ID | Forward primer | Reverse primer |
|---|---|---|
| **Mouse** | | |
| Hmg20a | AACCAACCCAGAGTTTGTGG | TTGCTCATCTTCAGGCCTTT |
| Gfap | GGGGCAAAAGCACCAAAGAAG | GGGACAACTTGTATTGTGAGCC |
| Vimentin | CGTCCACACGCACCTACAG | GGGGGATGAGGAATAGAGGCT |
| Il1b | AACTGTTGGTGAGGAATGTGG | GGTCCTGTCCCTCTTGTTTTCA |
| Slc1a2 | ACAATATGCCCAAGCAGGTAGA | CTTTGGCTCATCGGAGCTGA |
| Ereg | CTGCCTCTTGGGTCTTGACG | GCGGTACAGTTATCCTCGGATTC |
| Srebf2 | GCAGCAACGGGACCATTCT | CCCCATGACTAAGTCCTTCAACT |
| Igfbp3 | GACGACGTACATTGCCTCAG | GTCTTTTGTGCAAAATAAGGCATA |
| Tgfb1 | CTCCCGTGGCTTCTAGTGC | GCCTTAGTTTGGACAGGATCTG |
| Vcam1 | TTGGGAGCCTCAACGGTACT | GCAATCGTTTTGTATTCAGGGGA |
| Gapdh | CACCAACTGCTTAGCCCC | TCTTCTGGGTGGCAGTGATG |
| Cyclophilin | ATGGCAAATGCTGGACCAA | GCCATCCAGCCATTCAGTCT |
| **Human** | | |
| HMG20A | GCATGAAGATGAGCAACGAA | GCTCATTCATGAACCGAACA |
| ACTB | AAACTGGAACGGTGAAGGTG | GTGGCTTTTAGGATGGCAAG |
| HIST1H2AB | CGGTGCTTGAGTACCTGACC | TTCACTTTCCCTTGGCCTTA |

**TABLE S1**: List of primers used in this study.

| Primary antibodies | Host | Dilution | Supplier | Catalog number |
|---|---|---|---|---|
| HMG20A | Rabbit | 1:100 | Sigma-Aldrich | HPA008126 |
| Insulin | Mouse | 1:500 | Sigma-Aldrich | I2018 |
| Glucagon | Mouse | 1:200 | Sigma-Aldrich | A944 |
| Somatostatin | Goat | 1:100 | Santa Cruz Biotechnology | SC-7819 |
| Vimentin | Rat | 1:200 | Santa Cruz Biotechnology | SC-32322 |
| GFAP | Mouse | 1:500 | Sigma-Aldrich | G3893 |
| GPADH | Rabbit | 1:1000 | Cell Signaling Technology | 14C10 |
| **Secondary antibodies** | **Host** | **Dilution** | **Supplier** | **Catalog number** |
| Alexa fluor 568 goat anti-mouse | Goat | 1:800 | Thermo Fisher Scientific | A11004 |
| Alexa fluor 488 goat anti-rabbit | Goat | 1:800 | Thermo Fisher Scientific | A11008 |
| HRP anti-rabbit | Goat | 1:1000 | Sigma-Aldrich | AP307P |

**TABLE S2**: List of antibodies used in this study.

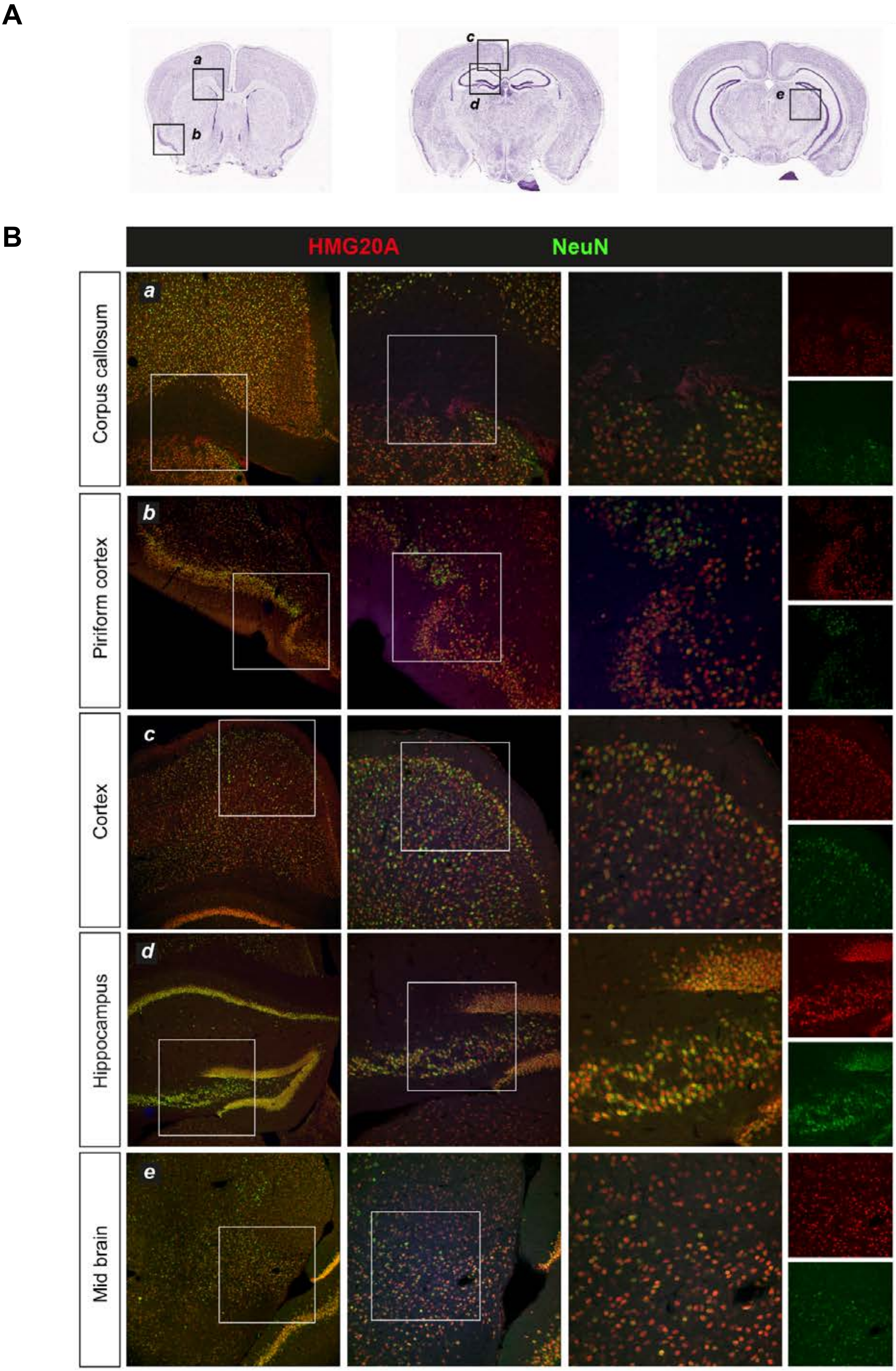


**Figure S1. HMG20A expression pattern in mouse brain astrocytes**. (**A**) Anatomical areas of the brain shown in (**B**) taken from the Allen Brain Atlas (https://mouse.brain-map.org/). (**B**) HMG20A (red) expression was assessed by immunofluorescence and was co-localized with NeuN (green, left panels) expressing astrocytes in various brain areas as depicted in the figure. Magnification: 10X

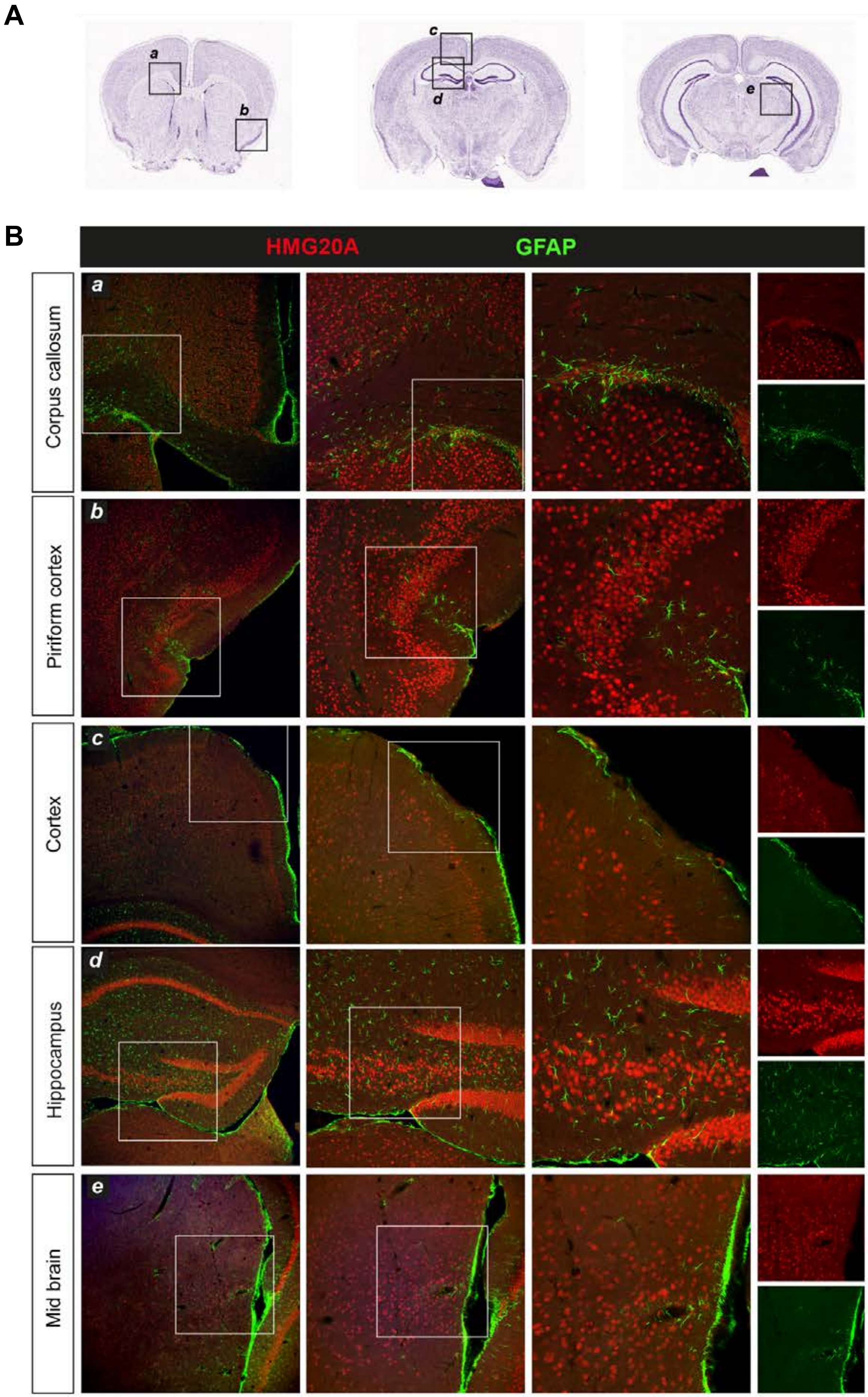


**Figure S2. HMG20A expression pattern in mouse brain neurons**. (**A**) Anatomical areas of the brain shown in (**B**) taken from the Allen Brain Atlas (https://mouse.brain-map.org/). (**B**)HMG20A (red) expression was assessed by immunofluorescence and was co-localized with NeuN (green, right panels) expressing neurons in various brain areas as depicted in the figure. Magnification: 10X

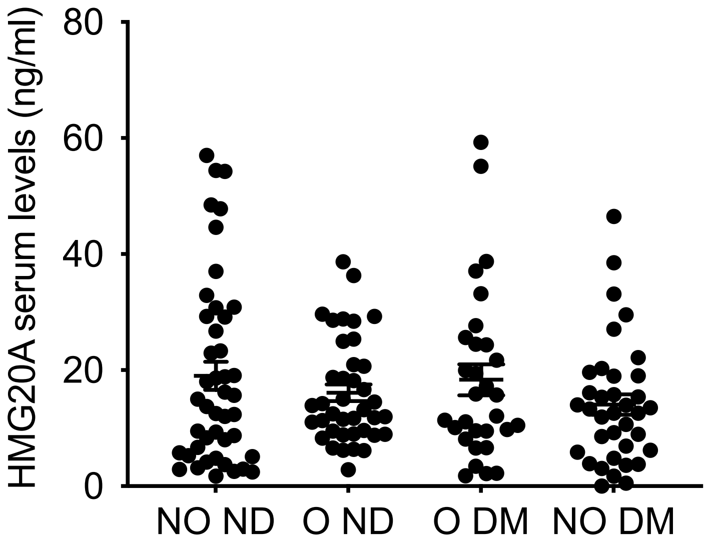


**Figure S3. HMG20A serum levels.** Quantification of HMG20A protein levels in serum isolated from the various groups depicted in the graph using an ELISA kit for human HMG20A. *n=40-50* individuals per group analyzed in duplicate.

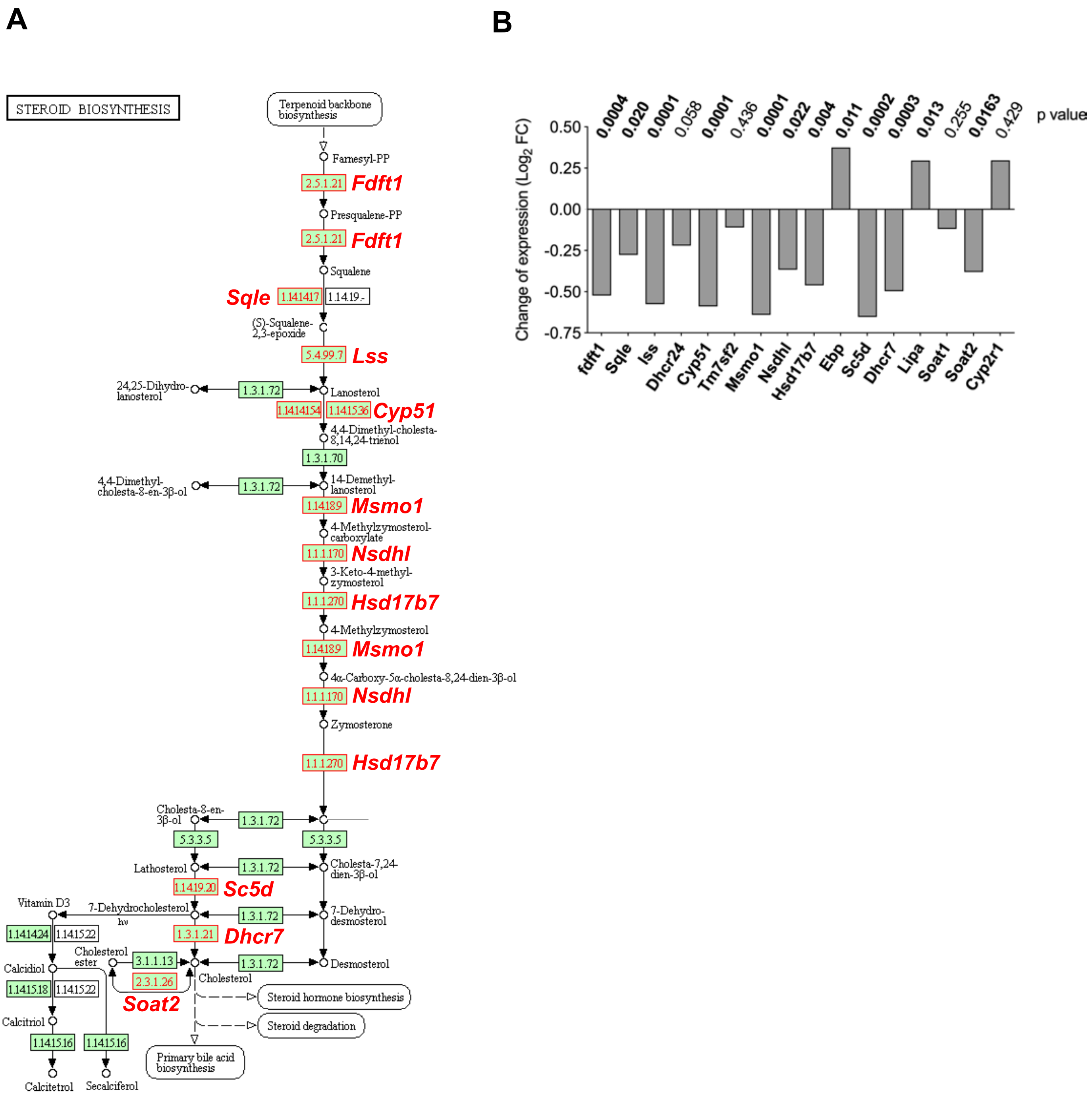


**Figure S4. Regulation of steroid biosynthesis pathway after *Hmg20a* silencing in astrocytes.** (**A**) **The** KEGG pathway of steroid biosynthesis showing the regulated genes after *Hmg20a* silencing. Enzymes present in mammals are highlighted in green. Enzymes whose encoding genes are downregulated after *Hmg20a* silencing are highlighted in red. *Mus musculus* gene names encoding the indicated enzyme are also included. (**B**) Bar Graph representing the relative expression levels of significantly regulated genes involved in sterol biosynthesis pathway, analyzed by LIMMA software. *p* value for each of the regulated genes is indicated above the bars. Values are referred to the expression levels detected in siCT.

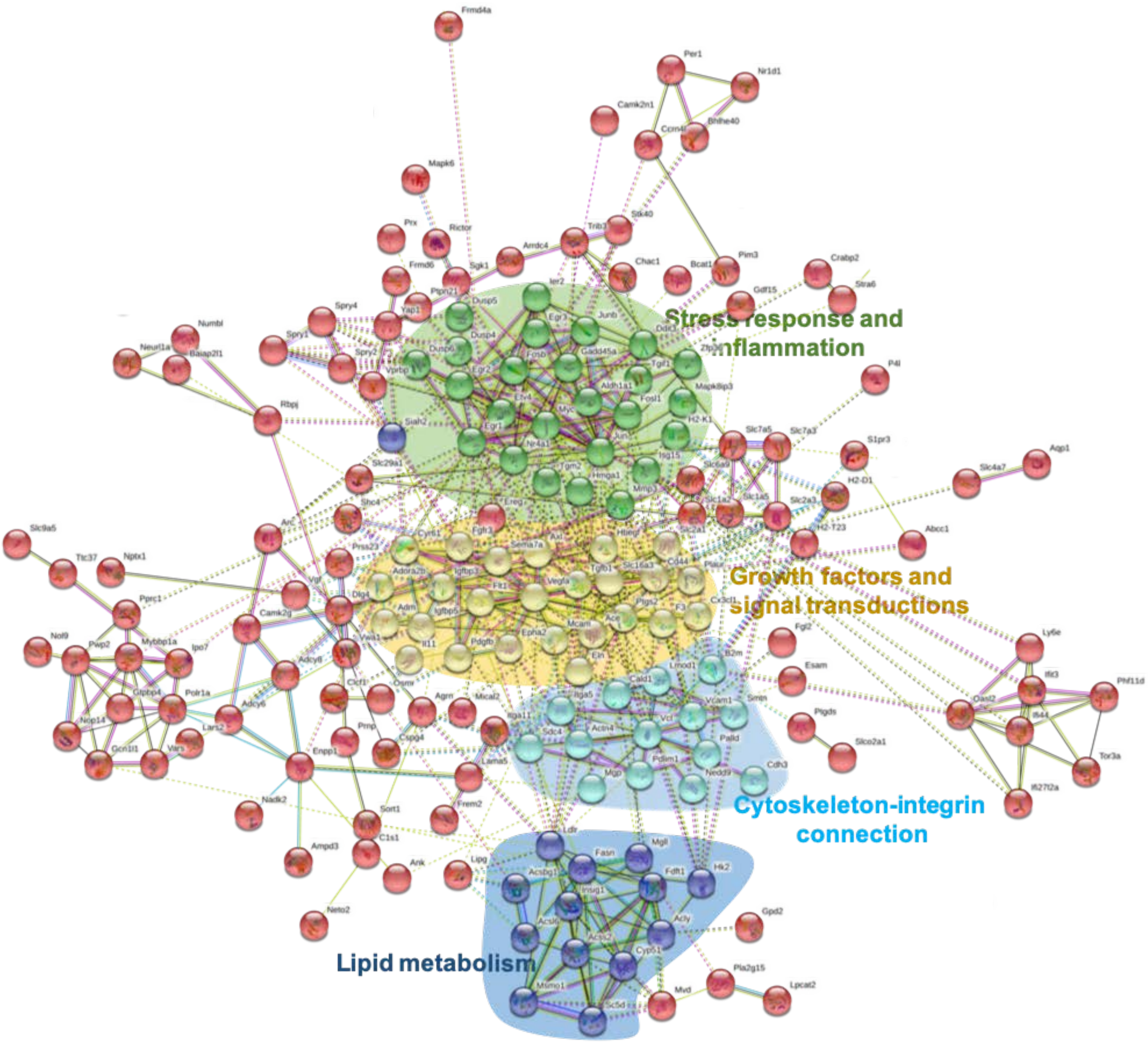


**Figure S5. Main biological networks regulated by HMG20A in astrocytes.** STRING cluster analysis of the 245 down-regulated genes in HMG20A repressed astrocytes. Only interacting mRNAs that were significantly repressed by siHMG20A are depicted. Criteria used were: Minimum required interaction score 0.4 and K-means clustering set to 5. The identified clusters are coloured and labelled as depicted in the figure. The solid and the dotted lines indicate connection within the same and different cluster respectively. Different color indicates different type of interactions: Cyan-from curated databases; Pink-experimentally determined; Blue-gene co-occurrence; Khaki-from text mining; Black-co-expression; Light blue-protein homology).

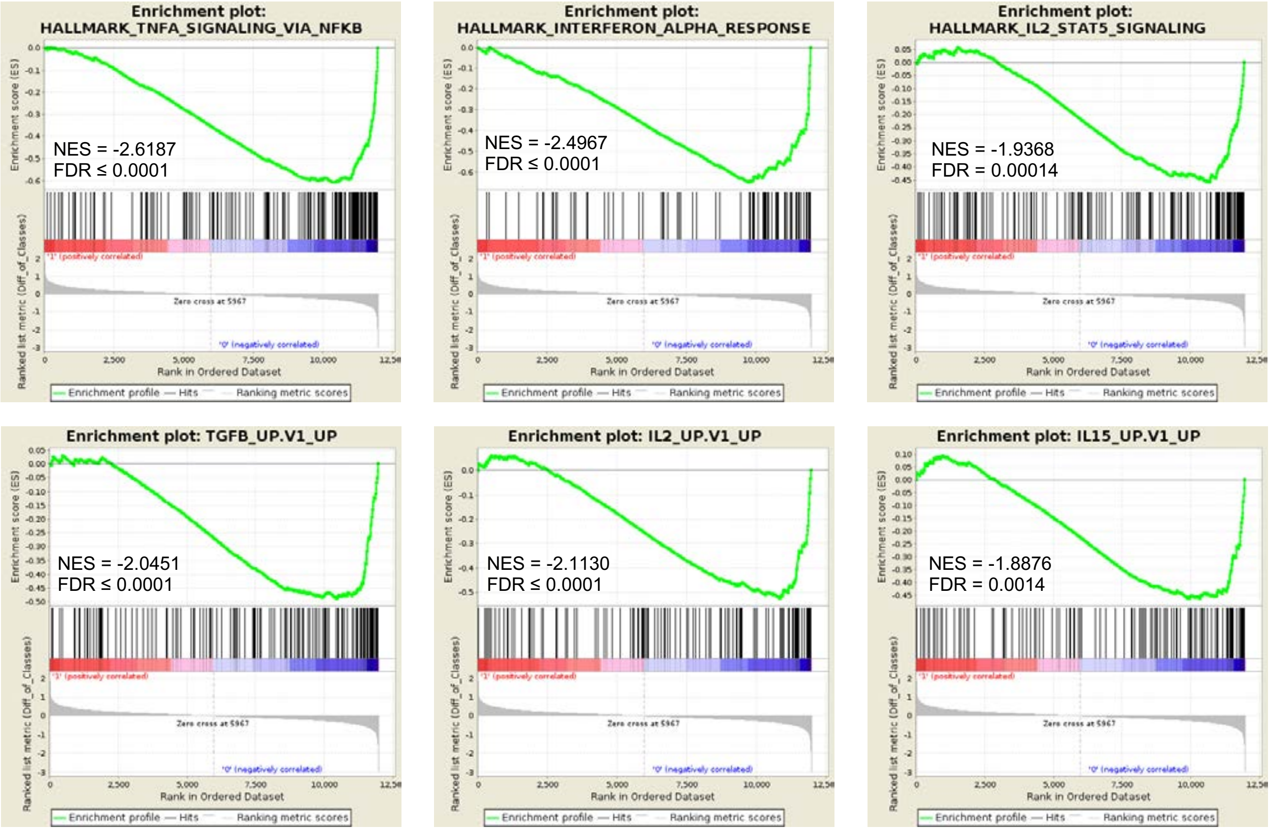


**Figure S6. Pathways enriched after *Hmg20a* silencing.** Enrichment plots of key inflammatory pathway enriched after *Hmg20a* silencing. The analysis was performed using the GSEA software. Normalized Enriched Score and FDR are provided within each plots.

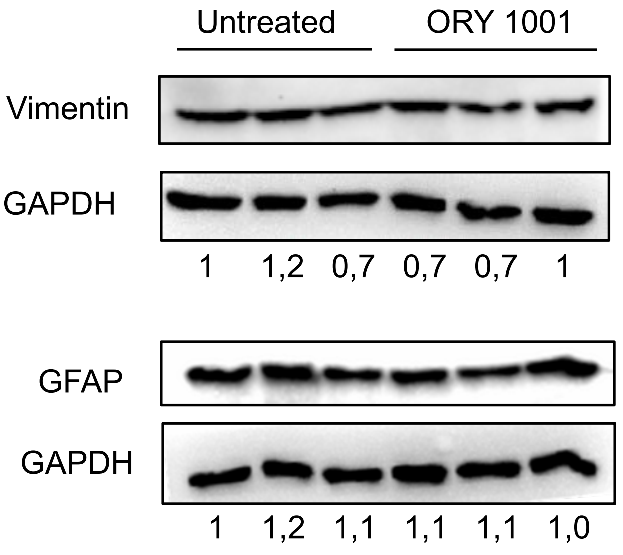


**Figure S7. GFAP and VIMENTIN levels in ORY-1001-treated astrocytes.** Protein levels of GFAP and VIMENTIN were assessed in whole cell extracts of astrocytes cultured or not with ORY-1001 for 72 hours. Values shown below are the relative expression levels of each protein normalized to GADPH.
